# Structural characterization of human tryptophan hydroxylase 2 reveals L-Phe as the superior regulatory domain ligand relevant for serotonin biosynthesis

**DOI:** 10.1101/2022.09.20.508773

**Authors:** Ida M. Vedel, Andreas Prestel, Zhenwei Zhang, Natalia T. Skawinska, Holger Stark, Pernille Harris, Birthe B. Kragelund, Günther H. J. Peters

## Abstract

Tryptophan hydroxylase 2 (TPH2) catalyzes the rate-limiting step in the biosynthesis of serotonin in the brain. Consequently, regulation of TPH2 is relevant for serotonin related diseases, yet, the regulatory mechanism of TPH2 is poorly understood and structural as well as dynamical insights are missing. Here, we use NMR spectroscopy to determine the structure of a 47 N-terminally truncated variant of the regulatory domain (RD) dimer of *human* TPH2 in complex with L-Phe, and show that L-Phe is the superior RD ligand compared to the natural substrate, L-Trp. Using cryo-EM we obtain a low-resolution structure of a similarly truncated variant of the complete tetrameric enzyme with dimerized RDs. The cryo-EM 2D class averages additionally indicate that the RDs are dynamic in the tetramer and likely exist in a monomer-dimer equilibrium. Our results provide structural information on the RD both as an isolated domain and in the TPH2 tetramer, which will facilitate future elucidation of TPH2’s regulatory mechanism affecting serotonin regulation.

## Introduction

Tryptophan hydroxylase (TPH, EC 1.14.16.4) is a non-heme, iron-dependent enzyme that catalyzes the first and rate-limiting step in the biosynthesis of the neurotransmitter and peripheral hormone, serotonin. TPH hydroxylates L-Trp into 5-hydroxy-L-Trp (5-HTP) using Fe(II) as co-factor and molecular oxygen and tetrahydrobiopterin (BH_4_) as co-substrates. 5-HTP is subsequently decarboxylated by aromatic amino acid decarboxylase to form serotonin^1^. There are two isoforms of TPH, independently regulating each their respective serotonin system ^2^: TPH1, mainly expressed in the gut and in the pineal gland, and TPH2, mainly expressed in the brain ^3^. Irregular levels of serotonin in the peripheral system (TPH1 associated) are related to e.g. irritable bowel syndrome and ulcerative colitis diseases ^4^, while irregular levels of serotonin in the brain (TPH2 associated) have been connected to e.g. schizophrenia ^5^ and major depression ^6^, highlighting TPH as a relevant drug target.

TPH constitutes together with phenylalanine hydroxylase (PAH, EC 1.14.16.1) and tyrosine hydroxylase (TH, EC 1.14.16.2) the family of aromatic amino acid hydroxylases (AAAH) ^7^. As TPH, PAH and TH catalyze the hydroxylation of their respective amino acid, L-Phe and L-Tyr. The AAAHs form homo-tetramers with each monomer consisting of an N-terminal regulatory domain (RD), a catalytic domain (CD), and a small tetramerization domain (TD, Figure 1A) ^8–11^. Of these, the RDs are the most divergent with relatively low sequence identity among the enzymes, while the CDs are very homologous ^12^. Between TPH1 and TPH2, the RDs are 48% identical (Figure 1A), and TPH2 contains an additional 46 N-terminal residues (N-terminal tail) in its RD associated with instability ^13,14^. The N-terminal tail contains an isoform specific phosphorylation site (Ser19) known to activate the enzyme upon phosphorylation and subsequent binding to 14-3-3 protein ^15^. The divergence between the RDs of TPH1 and TPH2 is promising in regards to selective targeting of one isoform over the other. However, to date there is no detailed structure available of either the RD of TPH2 or TPH1 and their exact regulatory mechanisms are yet to be determined. Therefore, insight into the RDs and their function is crucially needed, to not only enable selective drug targeting, but also to further elucidate how serotonin levels in the body are regulated.

**Figure 1.**
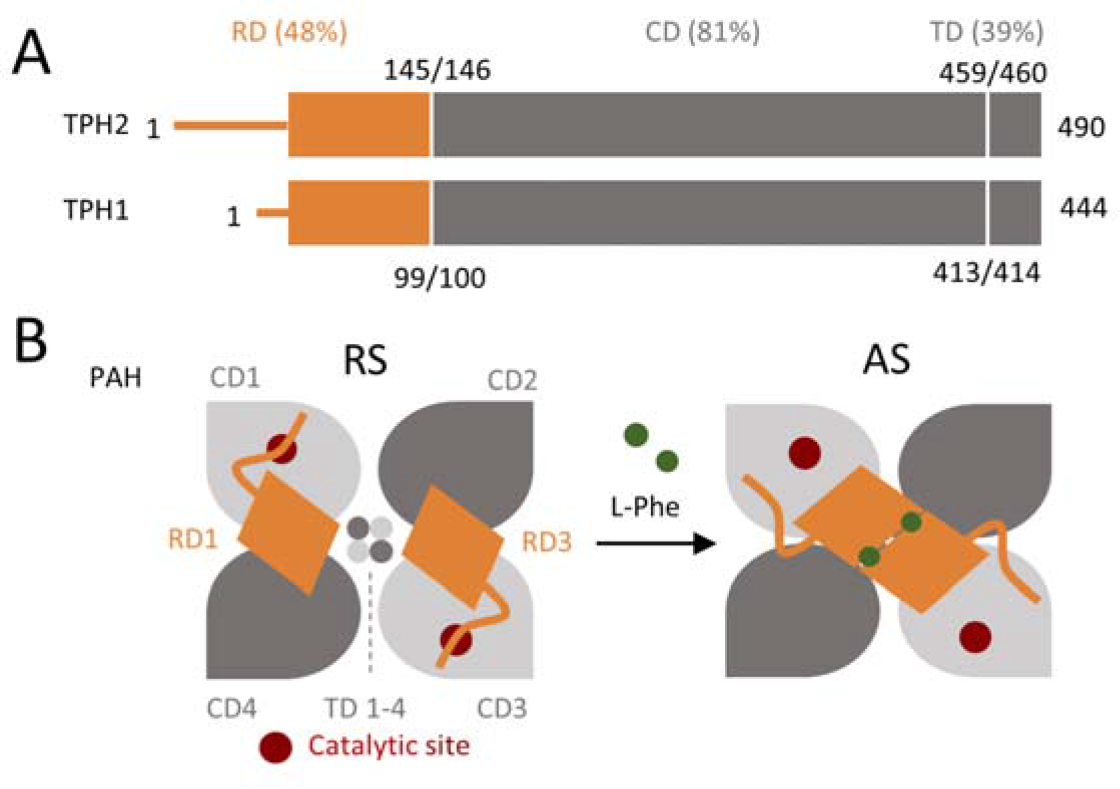
A) Domain alignment of human TPH2 and human TPH1 indicating the relative sizes of the regulatory domains (RDs), the catalytic domains (CDs), and the tetramerization domains (TDs). The sequence identities are indicated above and were determined using Clustal Omega ^65^, and numbers (in black) indicate which residue positions define the domain borders. Despite a low sequence identity, the TDs are expected to be structurally similar consisting of a single α-helix, which in the tetramer forms a 4-α-helix bundle with the other TDs ^7^ B) Illustration of the activation of PAH by binding of L-Phe, made with inspiration from ^20^. L-Phe shifts the conformation from the resting state (RS) to the active state (AS)^20^. The RS and AS have been confirmed by multiple crystal structures ^9,20,21^and small angle X-ray scattering (SAXS) data^20,23^.

Current knowledge on the RD of TPH2 is extrapolated from PAH as its regulatory mechanism is relatively well understood: PAH is allosterically activated by its substrate L-Phe, which has been confirmed by multiple kinetic studies ^16–19^. Structurally, the activation can be described by an equilibrium between a resting state (RS) and an active state (AS) with L-Phe bound ^20^, as shown in Figure 1B. In the RS, the RDs are docked between two CDs, and residues from the N-terminal tails hinders substrate access to the active site ^9,20,21^. In the AS, the RDs of two diagonally opposed monomers form a dimer with 2 L-Phe molecules bound at the dimer interface ^22^, resulting in the N-terminal tails moving away from the catalytic sites activating the enzyme ^20,23^. In solution, the isolated RD of PAH exists in an L-Phe affected monomer-dimer equilibrium ^22,24^. A similar equilibrium was observed by Tidemand et al., for an N-terminally truncated TPH2 variant containing the RD and CD (NΔ47-TPH2-RC) ^14^. Additionally, differential scanning fluorimetry (DSF) showed that both L-Phe and L-Trp have a stabilizing effect on the variant both indicating the presence of an RD binding site ^14^. This led to the hypothesis that TPH2 contains an allosteric site as PAH, but kinetic experiments have not revealed any substrate induced activation of TPH2 ^25,26^, indicating that TPH2 does not share the regulatory mechanism of PAH.

In this study, we structurally characterize the regulatory domain of *human* TPH2 and L-Phe binding to the domain. Using solution nuclear magnetic resonance (NMR) spectroscopy, we determine the structure of a 47 N-terminally truncated variant of the isolated RD (NΔ47-TPH2-R) in complex with L-Phe, and we observe that L-Phe binds stronger to NΔ47-TPH2-R than L-Trp. Furthermore, using cryo-electron microscopy (cryo-EM), we show that the RDs mainly form dimers in tetrameric TPH2 in a structure similar to the AS of PAH and additionally observe that the RDs are dynamic likely existing in a monomer-dimer equilibrium. Combined, these results establish L-Phe as the superior ligand of the isolated RD, and additionally provide a structural scaffold that will be important for studies of TPH2 regulation.

## Results

### The isolated regulatory domain of TPH2 is a dimer and L-Phe increases its thermal stability

To investigate if NΔ47-TPH2-R exists in a similar L-Phe-affected monomer-dimer equilibrium as NΔ47-TPH2-RC, analytical size exclusion chromatography was run at various concentrations of NΔ47-TPH2-R in the presence and absence of L-Phe (10 mM). L-Phe was used as the ligand since the natural substrate L-Trp interferes with the absorption at 280 nm. In the concentration range of 3-100 μM, NΔ47-TPH2-R eluted as a dimer for all protein loading concentrations and independent of the addition of L-Phe (Figure 2A-B). This is similar to the behavior of the isolated RD of TPH1, which also has been shown to exist as a dimer at protein concentrations above 2.7 μM ^27^. Although L-Phe did not affect the oligomeric state of NΔ47-TPH2-R, nanoDSF measurements showed that L-Phe had a stabilizing effect on the dimeric complex with an increase in melting temperature of up to 18 °C, consistent with the presence of a binding site as previously proposed ^14^ (Figure 2C). For the isolated RD of PAH (PAH-R) a similar increase in melting temperature of ~20 °C has been observed in the presence of 1 mM L-Phe ^22^.

**Figure 2.**
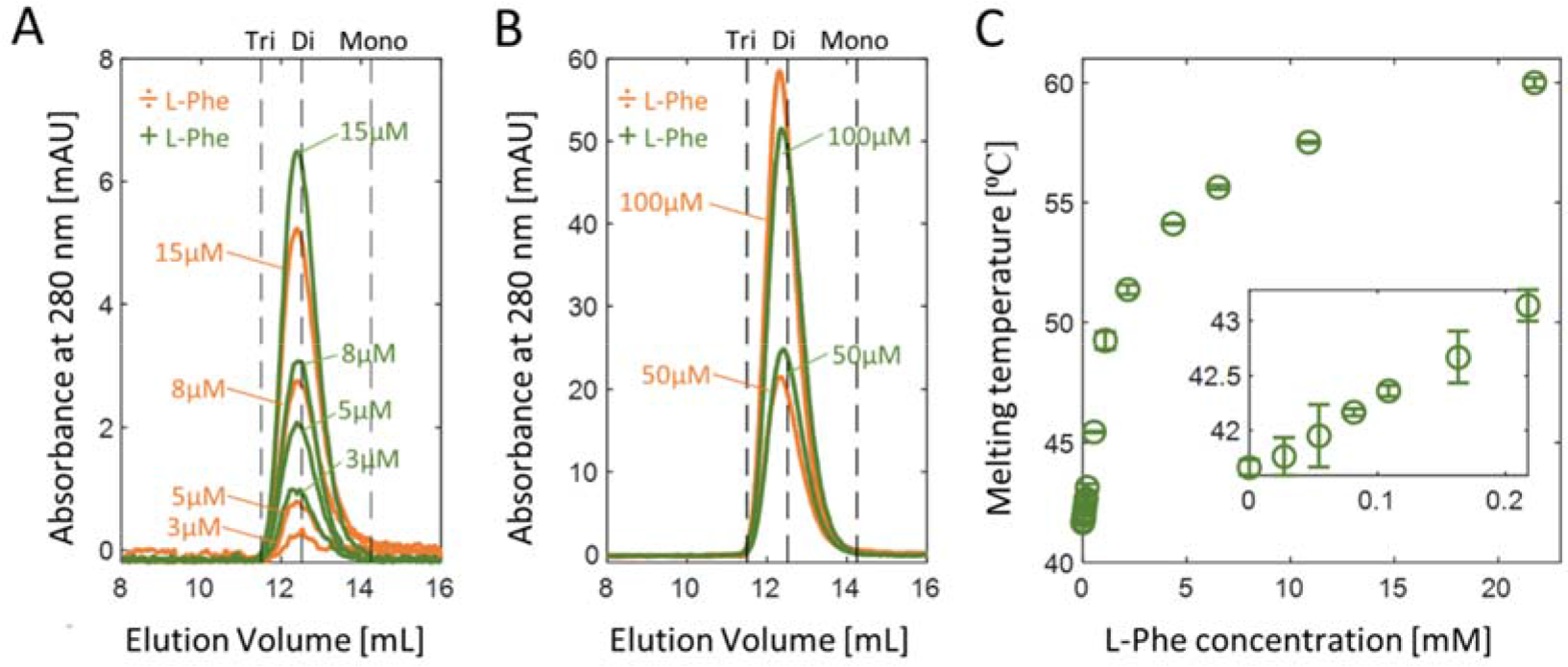
SEC runs with A) low protein concentrations (3-15 μM) and B) high protein concentrations (50-100 μM) with 0 mM L-Phe (orange) and with 10 mM L-Phe (green). The dashed lines indicate the expected elution volume of a monomer (Mono), dimer (Di), and trimer (Tri). C) The effect of L-Phe on the melting temperature of NΔ47-TPH2-R determined by nanoDSF. The mean of triplicate measurements (green circles) is shown with error bars representing the standard deviation. The insert shows a zoom-in of the low L-Phe concentration range.

### Solution structure of NΔ47-TPH2-R in complex with L-Phe reveals a binding site similar to the allosteric site of PAH

While free NΔ47-TPH2-R aggregated over time even at 4 °C, the presence of L-Phe in the buffer increased the stability and caused the domain to be stable at room temperature, which facilitated a detailed structural characterization. Here, we used solution NMR spectroscopy to obtain the first three-dimensional structure of the isolated RD of TPH2 in complex with L-Phe.

A ^1^H-^15^N HSQC spectrum of NΔ47-TPH2-R with L-Phe (Extended data figure 1A) showed 84 distinct backbone peaks out of the expected 96, confirming that the isolated RD forms symmetric dimers like the RDs of both PAH and TH ^22,28^. Additionally, an aromatic ^1^H-^13^C HSQC spectrum of ^13^C-labeled L-Phe and unlabeled NΔ47-TPH2-R, showed three peaks from the free L-Phe and three peaks from bound L-Phe, confirming that there is only one type of binding site (Extended data figure 1B). The quality of the L-Phe:NΔ47-TPH2-R NMR spectra enabled assignment of the majority of resonances of the complex and allowed for structure calculations. Inter- and intramolecular nuclear Overhauser effects (NOEs) were obtained from semi-automated NOE assignment providing 2171 distance restraints for the structure calculations (Extended data table 1). During the NOE assignment, P2 symmetry was imposed on both the NΔ47-TPH2-R dimer and the two L-Phe ligands. A total of 100 structures were calculated and refined with P2 symmetry imposed on the folded region, and the 20 lowest energy structures selected to represent the L-Phe:NΔ47-TPH2-R structure (Extended data table 1). The structure revealed that the RD of TPH2 contains a folded core with βαββα topology (β1: 65-71, α1: 76-85, β2: 90-98, β3: 105-112, α2: 116-129) and disordered N- and C-terminal tails (Figure 3A-B). The two monomers are arranged side-by-side forming an extended six stranded β-sheet. Although two residues (Thr134 and Leu135) show β-strand propensity based on their chemical shifts (Extended data figure 1C), a fourth β-strand is not fully formed (Figure 3B-C). Backbone ^15^N relaxation rates of L-Phe:NΔ47-TPH2-R support a folded core and flexible N- and C-terminal tails (Extended data figure 2). Additionally, modelling the flexible tails (residues 46-60 and 136-145) using SAXS data resulted in a very good fit (χ^2^=0.39, extended data figure 3 and extended data table 2), increasing the confidence in the NMR structure. The folded core confirms that the RD of TPH2 contains an ACT domain (aspartate kinase, chorismate mutase, and TyrA domain), a domain found in several proteins involved in amino acid metabolism including PAH and TH ^29^. The domain is characterized by βαββαβ topology, and the ACT domain often has an allosteric function in its parent enzyme by ligand binding to the domain ^29^. In some ACT domain enzymes, ligand binding is observed to cause large conformational changes as the binding of L-Phe to the RD of PAH, while more subtle allosteric changes occur in other ACT domain enzymes ^29,30^.

**Figure 3.**
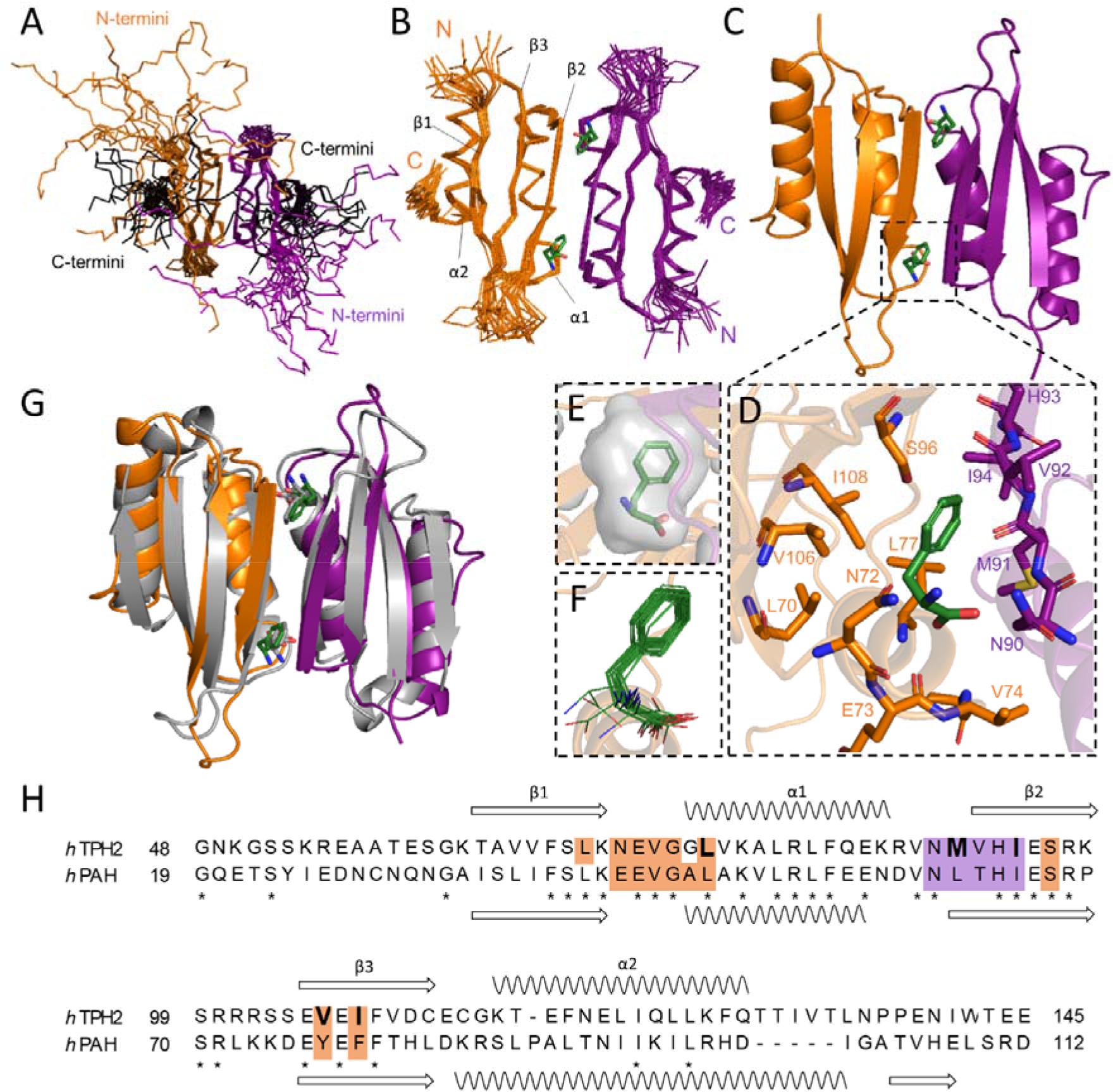
A) Superimposition of the 20 lowest energy conformers of the calculated NMR ensemble showing all residues (46-145) of monomer 1 (orange) and monomer 2 (purple). B) Superimposition of the 20 lowest energy conformers of the calculated NMR ensemble showing only the folded core (residues 61-135) of monomer 1 (orange) and monomer 2 (purple). L-Phe ligands are displayed as green sticks. C) Representative conformer of the calculated ensemble displayed as cartoon with monomer 1 (orange), monomer 2 (purple), and L-Phe ligands (green sticks). D) zoom-in on the residues in the binding pocket of the conformer in C. E) The void available in the binding pocket displayed as a grey blob (Illustrated using PyMOL 2.3.3 (Schrödinger), standard settings). F) Overlay of the position of L-Phe in the 20 lowest energy conformers. G) Superimposition of the conformer in C (orange/purple) with the crystal structure of human PAH-R (grey, PDB: 5FII ^22^). L-Phe bound in TPH2 (green sticks), L-Phe bound in PAH (grey sticks) H) sequence alignment between residues 48-145 of human TPH2 and residues 19-112 of human PAH. The secondary structure elements of NΔ47-TPH2-R are marked above the alignment. Residues involved in the binding of L-Phe in the dimer interface are marked with orange (monomer 1) and purple (monomer 2). Residues that showed cross peaks in the filtered ^13^C-NOESY-HSQC are highlighted in bold. * marks identical residues.

The two L-Phe ligands are bound at the dimer interface in hydrophobic pockets. Pocket 1 is formed by the side chains of Leu70, Leu77, Ser96, Val106, and Ile108 from monomer 1 and Met91 and Ile94 from monomer 2 (Figure 3D). Additionally, the backbone atoms of residues 72-75 (monomer 1) and residues 90-93 (monomer 2) are in close proximity to the ligand. The side chain of Asn72 and the backbone N-H of Leu77 are in positions to form hydrogen bonds to the amine group and the carboxyl group of L-Phe, respectively. Due to symmetry, pocket 2 is constituted by identical residues but from opposite monomers. The distance restraints for orienting L-Phe in the binding pockets were determined from a ^13^C/^15^N-filtered ^13^C-NOESY-HSQC spectrum, which exclusively yields NOESY cross peaks from ^13^C-labeled protein to unlabeled L-Phe. Five residues (Leu77, Met91, Ile94, Val106, Ile108) showed distinct cross peaks to the ligand (Extended data figure 4), establishing these residues as important parts of the L-Phe binding pockets. The binding pockets are small (160±40 Å^3^) with only little excess space available to the ligands, resulting in a tight fit (Figure 3E). The ligands are positioned very similarly in all 20 models, increasing the confidence in their position (Figure 3F).

A comparison of the structure of NΔ47-TPH2-R to the crystal structure of PAH-R (PDB: 5FII ^22^) showed a relatively high structural identity (RMSD: 2.1 Å), including the position of the L-Phe ligands (Figure 3G). A sequence alignment of the two regulatory domains (Figure 3H) showed that they are just 33% identical. However, the identical residues are clustered around the dimer interface where the binding pockets are located, a feature which has likely conserved the ability of TPH2 to bind L-Phe. Indeed, mapping the binding sites from each structure onto the sequence alignment showed that the binding pockets of L-Phe are located at almost the exact same place in the two enzymes, though the residues involved in binding are not completely conserved.

### L-Phe binds stronger to the regulatory domain of TPH2 than L-Trp

To investigate the binding of both L-Phe and L-Trp to the RD of TPH2, ^1^H-^15^N HSQC spectra of NΔ47-TPH2-R were recorded in the presence and absence of either L-Phe or L-Trp. The spectra show that the addition of 1 mM L-Phe causes large global changes to the spectrum, while addition of 20 mM L-Trp only causes very small chemical shift perturbations (CSPs) (Figure 4A-B). The gradual movement of the peaks in the L-Trp spectra from 0 mM-3 mM-20 mM shows that L-Trp binds in fast exchange and indicates a *K*_d_ in the mM range. In contrast, an NMR titration with L-Phe showed that L-Phe binds with slow exchange kinetics and that the complex is saturated at a concentration of 1 mM L-Phe (Extended data figure 5A-B). This reveals that L-Phe binds stronger to the RD dimer than L-Trp, despite L-Trp being the natural substrate of the enzyme. Due to the slow exchange kinetics, the assigned peaks from the L-Phe:NΔ47-TPH2-R complex could not be traced through the titration, and the spectrum of free NΔ47-TPH2-R was therefore assigned to enable calculation of CSPs and *K*_d_ determination by 2D NMR line-shape analysis. However, the aggregation propensity of free NΔ47-TPH2-R limited the feasible experimental conditions both in relation to temperature and protein concentration and thus resulted in low quality 3D spectra. Nonetheless, it was possible to assign 75 out of 93 peaks and CSPs were calculated and mapped onto the NMR structure (Figure 4C-F). For both L-Phe and L-Trp binding, almost all of the identified residues have some CSP, but with vast differences in amplitudes, where L-Phe binding causes around ten times larger CSPs than binding of L-Trp. It should be noted that the highest L-Trp concentration used was 20 mM, and it is possible that the L-Trp:NΔ47-TPH2-R complex is still not saturated. In both cases, the largest CSPs occur at the dimer interface where the binding pockets are located. For L-Phe binding there are especially large CSPs around loop 1 and loop 2 (L1 and L2), while L-Trp causes the largest CSPs around L2. It is possible that L-Trp is too large to fit properly into the binding pockets or that particular chemical complementarity is missing to anchor L-Trp in the pockets, giving rise to weaker binding and different CSP profile. Despite the large differences in the ^1^H-^15^N HSQC spectra of free and L-Phe bound NΔ47-TPH2-R, *R_1_* and *R_2_* backbone ^15^N relaxation rates and secondary structures based on secondary chemical shifts of the free and bound form are all similar, indicating that the overall size and shape of NΔ47-TPH2-R remains the same (Extended data figure 6).

**Figure 4.**
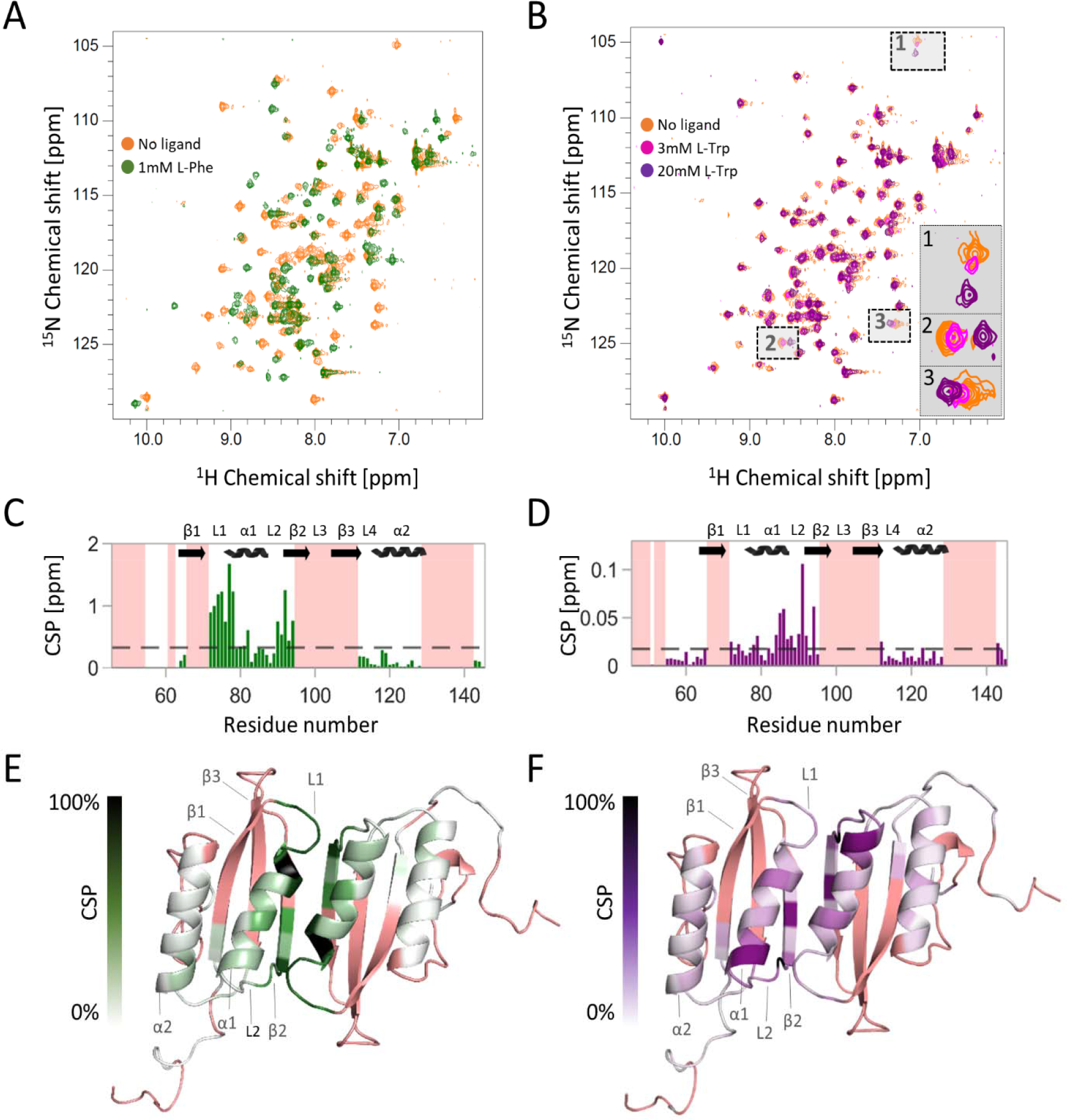
A) ^1^H-^15^N HSQC of NΔ47-TPH2-R at 25 °C in the presence and absence of L-Phe (1 mM) B) ^1^H-^15^N HSQC of NΔ47-TPH2-R at 25 °C in the presence and absence of L-Trp (3 mM and 20 mM). Inserts show zoom-in of areas 1, 2, and 3. C) Chemical shift perturbations caused by L-Phe binding at 25 °C. D) Chemical shift perturbations caused by 20 mM L-Trp binding at 25 °C. E) L-Phe perturbations mapped on the NΔ47-TPH2-R structure. F) L-Trp perturbations mapped on the NΔ47-TPH2-R structure. The red areas in C) and D) and the corresponding red coloring on the structures in E) and F) mark residues where it was not possible to calculate CSPs due to missing assignments.

The titration of NΔ47-TPH2-R with L-Phe, revealed peaks from two different states: The initial peaks from the free NΔ47-TPH2-R dimer and the final peaks from the NΔ47-TPH2-R dimer with two L-Phe molecules bound (Extended data figure 5B-C). Despite that a third state, the NΔ47-TPH2-R dimer with only one L-Phe bound was expected to exist, its population was too low to result in any observable peaks. Therefore, the titration data was fitted to a two state binding model considering the binding of one L-Phe molecule per NΔ47-TPH2-R monomer using the 2D line-shape analysis program TITAN ^31^ (Extended data figure 5C). The resulting *K*_d_ was 40.8±0.8 μM and *k*_off_ 0.02±0.05 s^−1^ (see Table 1). As the exchange rate is very slow, the *k*_off_ is at the limit of what can be reliably determined by line-shape analysis, and the *k*_off_ is therefore associated with a relatively large error. However, *k*_off_ is reliably small and likely to be less than 1 s^−1^. To increase the confidence in the determined *K*_d_, another titration was performed, by varying the concentration of unlabeled protein at a fixed concentration of ^13^C-labeled L-Phe. At low protein concentrations, the recorded aromatic ^1^H-^13^C HSQC spectra showed only three peaks arising from the free L-Phe. Only at the two highest protein concentrations, three peaks from bound L-Phe started to appear. Therefore, despite expected differences in *R*_2_ between bound and free L-Phe, a global fit was performed to the decreasing intensities of the three free L-Phe peaks (Extended data figure 7). The *K*_d_ was determined to 48±10 μM, which is in agreement with the one determined by line-shape analysis (40.8±0.8 μM), supporting a *K*_d_ in the 40-60 μM range.

**Table 1.** Calculated dissociation constants (K_d_) of the L-Phe:NΔ47-TPH2-R complex

| | $K_d$ ($\mu\text{M}$) | $k_{\text{off}}$ ($\text{s}^{-1}$) |
| --- | --- | --- |
| Protein observed titration (*line-shape analysis*) | $40.8 \pm 0.8$ | $0.02 \pm 0.05$ |
| Ligand observed titration (*Intensity fit*) | $48 \pm 10$ | NA |

### Cryo-EM images show dynamic regulatory domains in tetrameric TPH2

To address the structural behavior of the RD of TPH2 in a more native context, we next used cryo-EM to investigate an N-terminally truncated TPH2 variant containing the RD, the CD, and the TD (NΔ47-TPH2). Cryo-EM grids with this variant were set up both without L-Phe and, as large amounts of free L-Phe lower the image contrast, with 50 μM L-Phe, thus not reaching full saturation of the complex.

3D reconstruction of the NΔ47-TPH2 particles from the grids with L-Phe resulted in a tetrameric structure, with the four CDs and TDs creating an x-shaped planar core, and RDs from two diagonally opposed monomers forming dimers above and below the plane (Figure 5A). The dimeric conformation of the RDs is in agreement with a previous SAXS model of NΔ47-TPH2 in the presence of L-Phe (Skawinska et al., in preparation ^32^). While reconstruction of just the CDs and TDs reached 3.9 Å (Figure 5B and Extended data table 3), the complete structure was limited at 8.9 Å (ranging from 6-9 Å of the CDs to 15-20 Å of the RDs). However, our NMR structure of the RD dimer and the crystal structure of the CD and TD (PDB 4V06) could be well fitted into the density. The structure reveals a gap of ~10 Å between the RDs and CDs (Figure 5A, zoom in) likely causing a considerable degree of flexibility between the domains that could have contributed to the relatively low resolution. We observe that the RD dimer is oriented differently in the TPH2 tetramer compared to the cryo-EM structure of full-length TH ^33^, indicating structural differences between TPH2 and TH tetramers. (Extended data figure 8A). Assessing the 2D class averages of side view images (Figure 5C), revealed three different particle types: The most abundant, P2, with RD density on both sides of the CDs, P1, with density on one side, and P0, with no visible RD density. The missing RD density in P1 and P0 indicates that the RDs are dynamic, and likely move randomly relative to the CDs when they are not dimerized. The dynamic behavior of the RDs is consistent with the monomer-dimer equilibrium of NΔ47-TPH2-RC observed by Tidemand et al ^14^. Additionally, our observation is in agreement with a study from Zhu et al., who found that the RDs of an N-terminally tagged full-length TPH2 variant were too dynamic to be observed in their cryo-EM structure of the TPH2 tetramer ^34^. Comparing the 2D class averages from the grids without L-Phe (Figure 5D) to the 2D classes with 50 μM L-Phe, did not reveal any significant differences in the overall structure, (Figure 5C-D). The top view images confirm that NΔ47-TPH2 is a tetrameric complex both with and without L-Phe, and the side view images revealed the same three particle types (P2, P1, and P0) in similar abundances. Thus, either L-Phe does not affect the RDs in tetrameric TPH2, or 50 μM L-Phe is not enough to fully induce RD dimerization. In support of the latter, L-Phe induces dimerization of NΔ47-TPH2-RC ^14^, and stabilizes both NΔ47-TPH2-RC ^14^ and NΔ47-TPH2-R observed in this study. It is therefore likely that a higher L-Phe concentration would have an effect on RD dimerization. We additionally set up grids with full-length TPH2, in an attempt to get structural information on the N-terminal tail (residues 1-46). However, the 2D class averages were of low quality, likely due to the aggregation propensity of full-length TPH2, and thus it would not be possible to reach a resolution were the density of the tail could be clearly identified. The 2D classes still showed similar particle types (Extended data figure 8B) indicating that the N-terminal tail does not strongly affect the conformation of the RDs in tetrameric TPH2.

**Figure 5.**
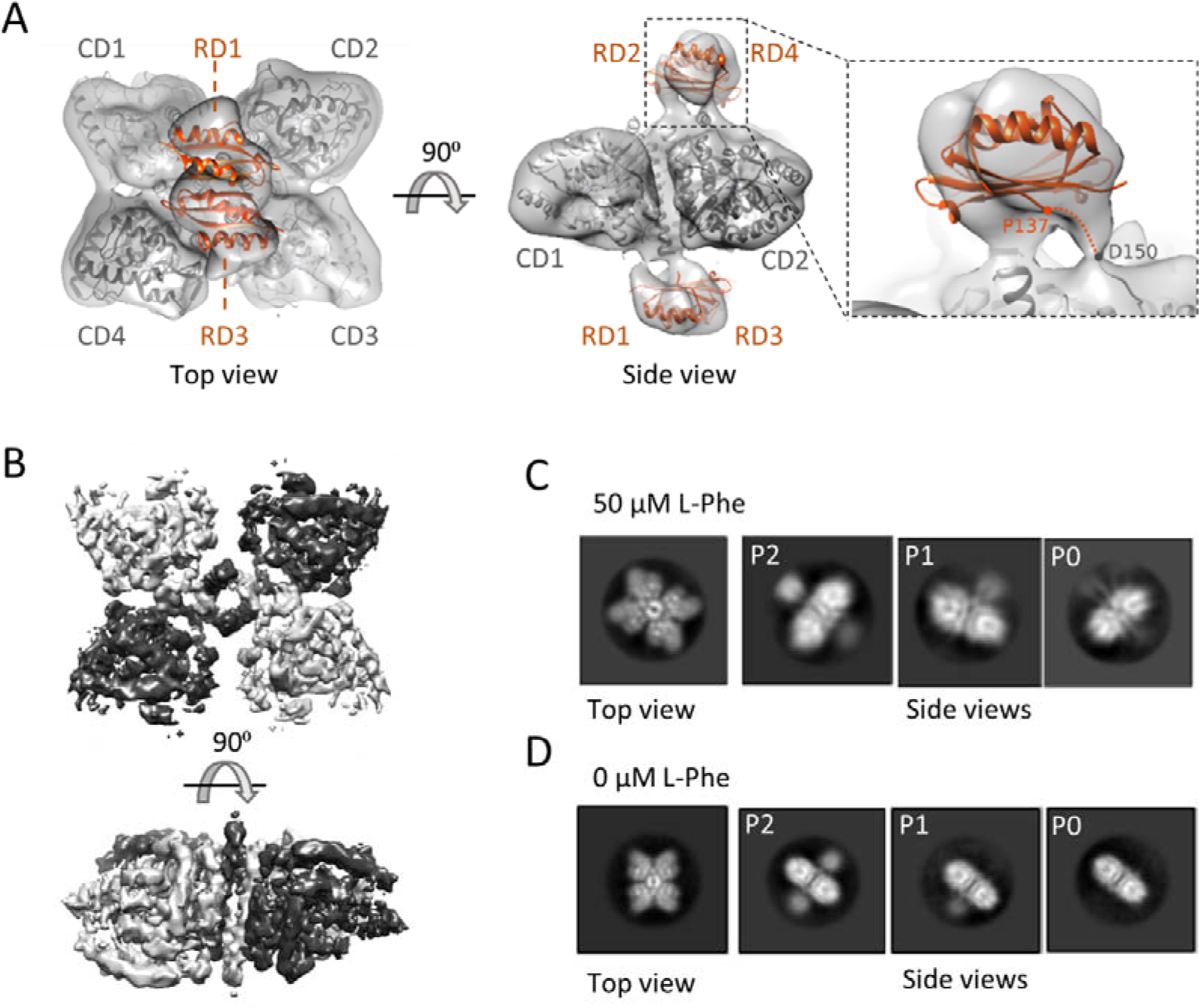
A) 3D reconstruction of NΔ47-TPH2 (Particle type P2) with 50 μM L-Phe. 8.9 Å EM map resolution. B) 3D reconstruction of the CDs and TDs of NΔ47-TPH2. 3.9 Å EM map resolution. C) Top view 2D class and side view 2D classes of the three different particle types of NΔ47-TPH2 with 50 μM L-Phe. D) Top view 2D class and side view 2D classes of the three different particle types of NΔ47-TPH2 without L-Phe.

## Discussion

In this work, we observe that NΔ47-TPH2-R exists as a dimer independent of the presence of L-Phe, whereas Tidemand et al. found that when the CD is included (NΔ47-TPH2-RC), the enzyme is in an L-Phe affected monomer-dimer equilibrium ^14^. Combined, this indicates that the presence of the CD weakens the RD dimer interface, highlighting the need for supplementary studies of full-length TPH2 to fully elucidate L-Phe binding to the RD. Nonetheless, the NMR structure of the L-Phe:NΔ47-TPH2-R complex presented in this study is the first structure of the isolated RD of TPH2, and it reveals the exact location of the RD binding sites, valuable for further characterization of the regulatory mechanism of TPH2. The binding sites are very similar to the allosteric sites of PAH, possibly resulting in L-Phe binding much stronger to the NΔ47-TPH2-R dimer than the natural substrate L-Trp. While the *K*_d_ of the L-Phe:NΔ47-TPH2-R complex is in the μM-range (40-60 μM), our NMR spectra indicate a *K*_d_ of the L-Trp:NΔ47-TPH2-R complex in the mM-range. The *K*_d_ of L-Phe:PAH-R has been determined to 8 μM ^24^ and 15 μM ^20^, and thus L-Phe binds slightly weaker to NΔ47-TPH2-R than to the isolated RD of PAH. The exact physiological levels of L-Phe in the brain of healthy individuals is, to the best of our knowledge, not known. However, Waisbren et al. found that elevated brain L-Phe levels of nine adults with early treated phenylketonuria (PKU, elevated L-Phe levels caused by a deficiency of PAH) ranged from 76-185 μM ^35^, whereas elevated brain L-Phe levels from 140-780 μM have been reported in other PKU patients ^36^. Thus, the *K*_d_ of L-Phe:NΔ47-TPH2-R reported in this study is within a physiological relevant range. Yet, we acknowledge that the *K*_d_ for both L-Phe and L-Trp may be different for full-length TPH2, where the monomer-dimer equilibrium of the RD is shifted, as observed in the cryo-EM images. Still, our results show that L-Phe is superior to L-Trp as a ligand for the isolated RD of TPH2.

Additionally, our cryo-EM data indicate that the RDs are dynamic in tetrameric TPH2, likely existing in a monomer-dimer equilibrium as illustrated in Figure 6, where RDs either form motionally restricted dimers or move randomly relative to the CDs as monomers in “undocked states”. The term undocked states was used by Khan et al. and Arturo et al. to describe the intermediate conformations between the RS and AS in PAH, where RD monomers are neither interacting with the CDs or another RD ^17,37^. Importantly, we do not observe any particles resembling the RS of PAH, but find a large part of the particles with dimerized RDs even without L-Phe in a conformation similar to the AS of PAH. Thus indicating that the conformational landscape of TPH2 is different to that of PAH. For PAH and TH, the N-terminal tail is creating an auto-inhibitory state by occluding the active site. While the N-tail of PAH is displaced by dimerization of the RDs induced by L-Phe binding ^20^, the N-tail of TH can reach and cover the active site with the RDs dimerized, and the tail is instead displaced upon phosphorylation of Ser40 ^33^. The N-tail of TPH2 is, similar to TH, longer than that of PAH. It is thus possible that residues in the end of the tail can cover the active site in a state with dimerized RDs, as also discussed by Zhang et al. ^27^. Although with low confidence, the Alpha-fold 2 predicts residues 1-21 of TPH2 to cover the entrance to the active site (Extended data figure 9) ^38^, suggesting at least that this region has a propensity to interact with the CD. This is consistent with activation by phosphorylation of Ser19 and 14-3-3 binding observed by Winge et al. ^15^, which could cause displacement of the N-tail. However, the binding of L-Phe to the RD dimer of TPH2 raises the question if TPH2 is also allosterically affected by L-Phe as PAH. Although we did not observe changes in the 2D classes of NΔ47-TPH2 in the presence of 50 μM L-Phe, the binding of L-Phe stabilized the RD (Figure 2C), and L-Phe affected the monomer-dimer equilibrium of TPH2-RC observed by Tidemand et al. ^14^, which jointly indicates that increasing concentrations of L-Phe would induce RD dimerization in tetrameric TPH2. L-Phe is already known to influence serotonin production to some extent, as reduced serotonin levels have been reported in PKU patients and is associated with some of the neurological symptoms of PKU ^39,40^. The mechanism of serotonin depletion in relation to PKU is not completely understood, but is hypothesized to be caused by either a lowered transport of L-Trp into the brain or increased competitive inhibition of L-Phe in the active site of TPH2 at elevated L-Phe levels ^39,40^, both resulting in reduced L-Trp hydroxylation. Thus, L-Phe binding to the RD could be an activation mechanism of TPH2, to keep a steady serotonin production regardless of minor increases in the L-Phe level. Though using TPH2 with an N-terminal MBP-tag expected to influence the N-tail, Ogawa et al. observed an inhibitory effect of L-Phe on TPH2 activity ^41^, indicating that L-Phe alone does not activate TPH2. Yet L-Phe RD binding could have a stabilizing effect, as also observed in this and in an earlier study ^14^, potentially influencing e.g. phosphorylation of the RD and subsequent 14-3-3 binding. However, with the current knowledge available on TPH2, the function of L-Phe binding to the RD remains speculative. Future experiments mapping the effect of L-Phe on TPH2 kinetics and phosphorylation will therefore be important to elucidate if L-Phe plays a role in serotonin regulation. Nonetheless, the current study has established L-Phe as a superior ligand of the RD of TPH2 compared to L-Trp, and shown that L-Phe affects the thermal stability of the RD dimer. Importantly, the study has provided detailed structural information on the RD of TPH2, which will be valuable in the elucidation of TPH2’s regulatory mechanism relevant for the understanding of serotonin related diseases.

**Figure 6.**
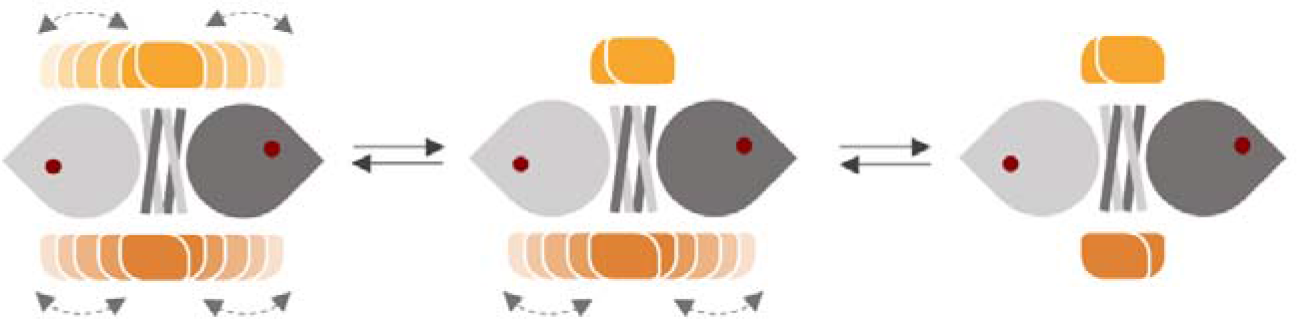
Schematic illustration of the proposed conformational landscape of TPH2, investigated using cryo-EM. The RDs (orange) can exist either as dynamic monomers randomly moving relative to the CDs (grey), or form more motionally restricted dimers.

## Materials and Methods

### Protein Expression

An LB agar plate with 30 μM/mL kanamycin sulfate (30 Kan) was streaked with a frozen stock of an *E. Coli* BL21 (DE3) strain containing the relevant MBP-tagged TPH2 construct (either MBP-NΔ47-TPH2-R, MBP-NΔ47-TPH2, or MBP-TPH2). The plate was incubated at 37 °C overnight. A single colony from the plate was used to inoculate 50 mL LB 30 Kan media in a 250 mL shaking flask and was incubated at 37 °C and 140 rpm shaking for around 4 h until the OD_600_ was 0.6-1.0. The cells were pelleted by centrifugation (1800 *g*, 4 °C, 10 min) and resuspended in 50 mL fresh LB 30 Kan media. 650 mL LB 30 Kan media in a 2 L shaking flask was then inoculated with 6.5 mL of the resuspended cells and incubated (30 °C, 140 rpm) for around 3h until the OD_600_ was 0.3. At this point, the temperature was lowered to 20 °C, the shaking set to 240 rpm, and the incubation continued for around 1.5 h until the OD_600_ was 0.6. Expression was induced by addition of isopropyl-β-D-thiogalactopyronoside (IPTG) to a final concentration of 0.1 mM. If the catalytic domain was part of the construct (MBP-NΔ47-TPH2 or MBP-TPH2), 650 μL of a sterile filtered freshly prepared (NH_4_)_2_Fe(SO_4_)_2_ solution was added to a final Fe(II) concentration of 0.2 mM. The culture was incubated for 16 h over-night (20 °C, 240 rpm) and cells harvested by centrifugation (4 °C, 3000 *g*, 15 min). The cells were resuspended in 20 mL specific binding buffer (Table 2), transferred to a 50 mL falcon tube, pelleted by centrifugation (4 °C, 3000 *g*, 15 min), and stored at −80 °C until purification. MBP-tagged 3C protease (MBP-3CP, for on-column cleavage of MBP-tagged TPH2) was expressed similarly, but incubated at 30 °C over-night. For expression of ^15^N- and ^15^N/^13^C-labeled MBP-NΔ47-TPH2-R for NMR experiments, the procedure was essentially the same but with the following modifications: two pre-cultures (50 mL LB 30 Kan media) were prepared and at OD_600_ 0.6-0.1, the cells were pelleted and resuspended in 20 mL 30 Kan M9 minimal media (22 mM KH_2_PO_4_, 53 mM Na_2_HPO_4_, 86 mM NaCl, 1 mM MgSO_4_, 1:1000 V/V M2 trace element solution, 22 mM glucose, 19 mM NH_4_Cl). If needed the glucose was ^13^C labeled and the NH_4_Cl ^15^N-labeled. The resuspended cells from both pre-cultures were used to inoculate 650 mL 30 Kan M9 minimal media in a 2L shaking flask and incubated as described above.

**Table 2.** Binding buffers used for purification of TPH2 variants

| TPH2 variant | Binding buffer |
| --- | --- |
| NΔ47-TPH2-R* | Std: 12 mM Na<sub>2</sub>HPO<sub>4</sub>/8 mM NaH<sub>2</sub>PO<sub>4</sub>, 100 mM (NH<sub>4</sub>)<sub>2</sub>SO<sub>4</sub>, pH 7.0 (adjusted with NH<sub>4</sub>OH) |
|  | L-Phe: 12 mM Na<sub>2</sub>HPO<sub>4</sub>/8 mM NaH<sub>2</sub>PO<sub>4</sub>, 100 mM (NH<sub>4</sub>)<sub>2</sub>SO<sub>4</sub>, 10 mM L-Phe pH 7.0 (adjusted with NH<sub>4</sub>OH) |
| NΔ47-TPH2<br>TPH2 | 20 mM Hepes, 300 mM (NH<sub>4</sub>)<sub>2</sub>SO<sub>4</sub>, pH 7.0 (adjusted with NH<sub>4</sub>OH) |

*All buffers were filtered using a 1000 mL filter unit with a 0.2 μm membrane (Fisher Scientific)*
*\* NΔ47-TPH2-R was purified in standard buffer (std) except for protein purified for NMR structure determination and SAXS experiments where the L-Phe buffer was used.*

### Protein purification

To purify the NΔ47-TPH2-R, frozen cells from a 650 mL culture containing MBP-NΔ47-TPH2-R were thawed and re-suspended in 40 mL specific binding buffer (Table 2). The cells were lysed while kept in ice-water using a Labsonic P sonicator (Sartorius), with 80% amplitude and 1 cycle for 3×30 s with 1 min cooling in-between. The sample was centrifuged (2×20 min, 4 °C, 16000 *g*), and the supernatant recovered and filtered through a 0.45 μm syringe filter (Whatman™). The supernatant was loaded onto a Dextrin Sepharose affinity column (GE Healthcare) pre-equilibrated with 5 CV of binding buffer, using an ÂKTA start chromatography system (GE Healthcare). The flow rate was 5 mL/min, and the progress of the purification was followed by UV absorbance at 280 nm. After loading the sample, un-bound proteins were removed with 5-6 CV of binding buffer. Hereafter, 30 mL of 2.2 μM MBP-3CP (diluted from a stock into binding buffer) was loaded on the column, and the system was paused for 1 h to ensure sufficient cleavage, leaving two additional residues from the linker on the N-terminal of the protein (Gly-Pro). After cleavage, NΔ47-TPH2-R was eluted from the column using binding buffer and ~8 mL protein collected. Elution buffer (binding buffer + 1 mM Maltose) was run through the column to elute MBP-3CP and cleaved MBP. The collected 8 mL protein was filtered through a 0.45 μm syringe filter (PALL), and loaded onto a HiLoad 26/600 Superdex 75 prep grade column (GE Healthcare), pre-equilibrated with 2 CV of binding buffer and with a flow rate of 2.6 mL/min. NΔ47-TPH2-R was collected from the peak eluting at the expected elution volume (confirmed by SDS-PAGE). The protein was concentrated to the desired concentration using a Vivaspin Turbo centrifugal concentrator (Sartorius) with a 3 kDa membrane by centrifugation at 4000 *g* and 4 °C. The protein was aliquoted, flash frozen in liquid N_2,_ and stored at −80 °C until further use. The purity was assessed by SDS-PAGE.

To purify NΔ47-TPH2 and TPH2 for use in cryo-EM experiments, the following modifications were done: The chromatographic system was an ÂKTA Pure (GE Healthcare) and a very narrow fraction of the top of the affinity chromatography peak after protease cleavage was collected. The concentration was measured on a NanoDrop One Microvolume spectrophotometer (Thermo Scientific™) and 1.5x molar equivalent of (NH_4_)_2_Fe(SO_4_)_2_ was added from a freshly prepared stock solution and allowed to incubate for around 15 min before filtering the sample through a 0.22 μm centrifuge tube filter (Corning). The protein was immediately used to set up cryo-EM grids. The purity was assessed by SDS-PAGE and negative staining.

MBP-3CP was purified similarly to NΔ47-TPH2-R except that the binding buffer was 20 mM Tris/HCl, 70 mM (NH_4_)_2_SO_4,_ pH 8.0 and that MBP-3CP was eluted directly using elution buffer (binding buffer + 0.1 mM maltose) after removing un-bound proteins with binding buffer. The buffer was then exchanged to 50 mM Tris/HCl, 50 mM (NH_4_)_2_SO_4,_ 10 mM EDTA, 1 mM DTT, 20 % V/V glycerol, pH 8.0 using an Amicon stirred cell with a PLGC NMWL 10 kDa membrane (EMD Millipore) and pressurized with N_2_, ensuring >500x dilution of the buffer. The MBP-3CP was aliquoted into stocks containing MBP-3CP for one TPH purification (80 nmol, ~0.5-1 mL), and stored in Eppendorff tubes at −20 °C.

### Sample preparation

If not stated otherwise, frozen samples of NΔ47-TPH2-R were prepared for experiments in the following way: A frozen sample of the relevant variant was thawed on ice and filtered through a 0.22 μm centrifuge tube filter (Corning). The protein concentration was determined as a mean of triplicate measurements using an ND-1000 NanoDrop Spectrophotometer (Saveen Werner). The extinction coefficient of NΔ47-TPH2-R at 280 nm were estimated based on the protein sequence using ProtParam from the ExPASy server ^42^.

### Analytical size exclusion chromatography (SEC)

The oligomeric state of NΔ47-TPH2-R was investigated by analytical size exclusion chromatography using a Superdex 75 10/300 GL column (GE Healthcare) and an ÂKTA Explorer chromatographic system (GE Healthcare). The expected elution volumes of a monomer, dimer, and trimer were estimated based on a calibration curve constructed from the column manual (Superdex 75 10/300 GL, GE Healthcare). The column had been pre-equilibrated with 2 CV of running buffer (Std buffer or L-Phe buffer, Table 2). Samples of 500 μL with the desired protein concentrations were prepared by diluting NΔ47-TPH2-R with either Std or L-Phe buffer. The samples were loaded using a 500 μL sample loop and eluted with 1.5 CV of running buffer at a flow rate of 0.8 mL/min.

### Nano differential scanning fluorimetry (nanoDSF)

The influence of L-Phe on the thermal stability of NΔ47-TPH2-R was measured using nanoDSF. 15 μL samples with a final concentration of 100 μM NΔ47-TPH2-R were prepared by mixing the relevant volumes of NΔ47-TPH2-R, (Std buffer Table 2), and an L-Phe stock solution (1.1 mM L-Phe for low concentrations/50 mM L-Phe for high concentrations in Std buffer). The measurements were performed with a Promotheus NT.48 (NanoTemper Technologies) using standard grade capillaries (NanoTemper Technologies) and run with a constant temperature ramp of 0.5 °C/min from 20 °C to 95 °C. All measurements were run in triplicates. The maximum of the first derivative of the thermal unfolding curves (350 nm/330 nm ratio) was used to determine the melting temperature of the individual measurements. The final melting temperature was calculated as a mean of the three measurements with standard deviation using Matlab (Mathworks Inc).

### Nuclear Magnetic Resonance (NMR) spectroscopy

NMR spectra were acquired on a Bruker Avance III 600 MHz or 750 MHz spectrometer with TCI cryo-probes. Spectra for structure determination, titrations, and backbone relaxation parameters were referenced internally to DSS at 0 ppm in the ^1^H dimension and by relative gyromagnetic ratios in the ^13^C and ^15^N dimensions. All data sets were processed and phase corrected manually with either Topspin 3.6 for 2D spectra or qMDD ^43^ and NMRPipe ^44^ for non-uniformly sampled 3D spectra and pseudo 3D spectra. The buffer conditions of NMR samples were 12 mM Na_2_HPO_4_/8 mM NaH_2_PO_4_, 100 mM (NH_4_)_2_SO_4_, pH 7.0 (adjusted with NH_4_OH), 125 μM DSS and 5% (v/v) D_2_O unless specified otherwise. Secondary structure propensities were calculated based on chemical shifts using Talos-N ^45^, and secondary ^13^C^α^ chemical shifts were calculated by use of random coil shifts estimated using the online tool from Kjaergaard et al. ^46^. Chemical shift perturbations were calculated using the following equation:

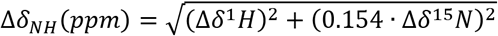

### NMR assignment

Resonances of free and L-Phe bound NΔ47-TPH2-R were assigned manually using CcpNmr analysis 2.4.2 ^47^. Backbone and sidechain resonances of L-Phe bound NΔ47-TPH2-R were assigned using following triple resonance spectra all measured at 30 °C: BEST TROSY (BT)-HNCA, BT-HNCACB, BT-HN(CO)CACB, BT-HNCO, BT-HN(CA)CO, HcCH-TOCSY, hCCH-TOCSY, ^15^N-NOESY-HSQC (120 ms mixing time), and ^13^C-NOESY-HSQC (120 ms mixing time) and BT-HNCA (measured at 40 °C). Additionally, three versions of amino acid specific ^1^H-^13^C HSQC spectra were used in the assignment process of the methyl resonances. The spectra were measured on one sample (A1) containing 1.1 mM NΔ47-TPH2-R and 10 mM L-Phe. Chemical shifts for the L-Phe ligands were obtained from a ^1^H-^13^C HSQC spectrum and an aromatic ^1^H-^13^C HSQC spectrum on a sample (A2) containing 450 μM unlabeled NΔ47-TPH2-R and 2.6 mM ^13^C/^15^N-labeled L-Phe. Backbone resonances of free NΔ47-TPH2-R were assigned using following triple resonance spectra collected on multiple samples: BT-HNCA (25 °C), BT-HNCO (25 °C), BT-HN(CO)CA (25 °C), BT-HNCACB (25 °C), HcCH-TOCSY (20 °C), hCCH-TOCSY (20 °C), BT-HN(CA)CO (20 °C), BT-HNCACB (20 °C), ^15^N-NOESY-HSQC (20 °C, 120 ms mixing time), ^13^C-NOESY-HSQC (20 °C, 120 ms mixing time). Additionally, an HNCA spectrum (20 °C) recorded using a gradient optimized CO decoupling pulse (GOODCOP ^48^), and three HNCA spectra (20 °C) with different variations of a beta/alpha decoupling pulse (BADCOP1, BADCOP2, BADCOP3 ^48^) were used in the assignment process. Details on the samples used are provided in Table 3.

**Table 3.** NMR samples of free NΔ47-TPH2-R

| Sample | NΔ47-TPH2-R<br>concentration* | Spectra |
| --- | --- | --- |
| B1 | 1000 μM | BT-HNCA (25 °C) |
| B2 | 200 μM | BT-HNCO (25 °C), BT-HNCOCA (25 °C) |
| B3 | 275 μM | BT-HNCACB (25 °C), HcCH-TOCSY (20 °C) |
| B4 | 250 μM | hCCH-TOCSY (20 °C), BT-HN(CA)CO (20 °C) |
| B5 | 240 μM | <sup>13</sup>C-NOESY-HSQC (20 °C), <sup>15</sup>N-NOESY-HSQC (20 °C) |
| B6 | 800 μM | BT-HNCACB (20 °C), BADCOP1 (20 °C),<br>BADCOP2 (20 °C), BADCOP3 (20 °C), GOODCOP<br>(20 °C) |

*\*The stated concentration is the initial concentration of the sample. Due to aggregation of the protein, the concentration gradually decreased over time. <sup>1</sup>H-<sup>15</sup>N HSQC spectra were recorded before and after each of the 3D spectra to ensure that the aggregation had not caused any structural changes in the remaining soluble protein.*

### NMR Structure calculations and refinements

Intra- and intermolecular (protein – protein) restraints for the structure calculations of NΔ47-TPH2-R in complex with L-Phe were obtained from a ^15^N-NOESY-HSQC spectrum and a ^13^C-NOESY-HSQC spectrum with 120 ms mixing time recorded on sample A1. Intermolecular restraints (protein – ligand) were obtained from a ^13^C/^15^N-filtered ^13^C-NOESY-HSQC and a ^13^C/^15^N -filtered ^15^N-NOESY-HSQC with 120 ms mixing time also recorded on sample A1. In all cases, NOE peaks were picked manually. Backbone dihedral restraints were calculated from the chemical shifts using TALOS-N ^45^. Using CYANA 3.98.5 ^49^, the NOESY spectra were semi-automatically assigned and initial structures were calculated to generate distance restraints. This process was done iteratively, and assignments and structures were evaluated manually and recalculated with modified inputs in each iteration. To unambiguously assign the peaks of the ^13^C/^15^N-filtered ^13^C-NOESY-HSQC spectrum and the ^15^N-filtered ^13^C/^15^N-NOESY-HSQC spectrum to the L-Phe ligands, the chemical shifts in the indirect ^1^H dimension of the filtered NOESY spectra and the ^1^H chemical shifts of the L-Phe ligands were shifted with +12 ppm. P2 symmetry were imposed on both the NΔ47-TPH2-R dimer and the two L-Phe ligands. XPLOR-NIH v2.44 ^50,51^ was subsequently used to calculate and refine 100 structures with the distance restraints generated from the initial structure calculations. The structures were refined using an implicit water potential (EEFx ^52^), and an energy term was added to restrain the folded regions (residues 63-135) in P2 symmetry. The 20 lowest energy structures were selected to represent the dimer structure of NΔ47-TPH2-R in complex with L-Phe.

### *NMR titrations and K*_d_ *determination*

For the titration of ^15^N-labeled NΔ47-TPH2-R with L-Phe, the concentration of NΔ47-TPH2-R was kept constant at 200 μM and ^1^H-^15^N HSQC spectra of the following NΔ47-TPH2-R:L-Phe molar ratios were recorded at 20 °C: 1:0, 1:0.3, 1:0.6, 1:0.9, 1:1.2, 1:1.5, 1:1.8, 1:2.2, 1:2.8, 1:4.2, 1:6. For lineshape analysis, the spectra were processed using an exponential window function with 4 Hz and 8 Hz line broadening in the ^15^N and ^1^H dimensions, respectively. The data was fitted to a 2-state binding model using the 2D lineshape analysis software TITAN ^31^. 21 peak pairs were used for the analysis and the errors were determined by a bootstrap analysis included in the TITAN software using 100 replicas.

For the titration of ^13^C-labeled L-Phe with NΔ47-TPH2-R the concentration of L-Phe was kept constant at 260 μM and aromatic ^1^H-^13^C HSQC spectra of the following L-Phe:NΔ47-TPH2-R molar ratios were recorded at 20 °C: 1:0, 1:0.06, 1:0.11, 1:0.23, 1:0.29, 1:0.34, 1:0.38, 1:0.46, 1:0.57, 1:0.76, 1:2.0. The spectra were processed using a QSINE window function with 1 Hz and 0.3 Hz line broadening in the ^1^H and ^13^C dimensions, respectively. Though differences in *R*_2_ between bound and free L-Phe is expected ^53^, the intensities of the three decreasing peaks were extracted using CcpNmr analysis and fitted to the following equation using Origin2019 (OriginLabs) with a shared *K*_d_ parameter for all three peaks.

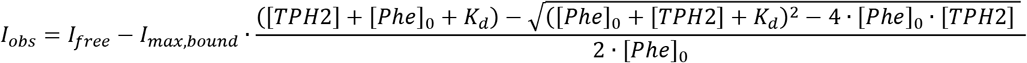

Where *I_obs_* is the intensity of a free L-Phe peak at a certain concentration of NΔ47-TPH2-R, *I_free_* is the intensity of a free L-Phe peak at 0 μM NΔ47-TPH2-R, *I_max,bound_* is the difference between *I_free_* and *I_obs_* at complete saturation, [*TPH*2] is the concentration of NΔ47-TPH2-R, [*Phe*]_0_ is the initial concentration of free L-Phe, and **K*_d_* is the dissociation constant.

### Backbone ^15^N relaxation parameters

Longitudinal (*R*_1_) relaxation rates, transverse (R_2_) relaxation rates, and steady state heteronuclear ^1^H-^15^N NOE relaxations were determined for L-Phe bound NΔ47-TPH2-R at 30 °C. Data sets were recorded on a sample containing 750 μM ^13^C/^15^N labeled NΔ47-TPH2-R and 10 mM L-Phe on both a 600 MHz and a 750 MHz spectrometer. A single data set was recorded for both the determination of *R*_1_ and *R*_2_ relaxation rates, while data sets for ^1^H-^15^N NOEs were recorded in triplicates on the 600 MHz spectrometer and duplicates on the 750 MHz spectrometer. To determine *R*_1_ relaxation rates, the experimental delay times were: 0, 100, 200, 400, 600, 900, and 1300 ms and for *R*_2_ relaxation rates the delay times were: 0, 8.48, 16.96, 33.92, 50.88, 76.32, 101.76, and 135.68 ms. The intensity of the spectra as a function of delay times were fitted to an exponential function using CcpNmr Analysis v2.5.1 ^47^. ^1^H-^15^N NOEs were calculated from the ratio of spectra with and without ^1^H pre-saturation and each data set was analyzed individually with CcpNmr Analysis v2.5.1 ^47^. The final ^1^H-^15^N NOEs and errors were calculated as a mean of the individually determined values with a standard deviation using Matlab (Mathworks Inc.).

*R*_1_ and *R*_2_ relaxation rates were determined for both free NΔ47-TPH2-R (150 μM ^15^N labeled NΔ47-TPH2-R) and L-Phe bound NΔ47-TPH2-R (150 μM ^15^N labeled NΔ47-TPH2-R and 10 mM L-Phe) on a 750 MHz spectrometer at 20 °C. All data sets were recorded in duplicates. To determine *R*_1_ relaxation rates, the experimental delay times were: 0, 100, 200, 400, 600, 900, and 1300 ms and for *R*_2_ relaxation rates the delay times were: 0, 8.48, 16.96, 33.92, 50.88, 76.32, 101.76, and 135.68 ms. The peak intensity with respect to delay time was fitted to an exponential function using CcpNmr Analysis 2.5.1 ^47^.

### L-Phe pocket volume calculations

The volume of both L-Phe pockets were calculated for the five lowest energy structures of the NMR ensemble using SiteMap (Schrödinger, Inc.). The binding site including 3Å buffer region was evaluated, and the rest of the settings were kept at default values. The final volume was calculated as the mean of all ten calculations with standard deviation using Matlab (Mathworks Inc).

### Small angle X-ray scattering (SAXS) data collection

Different concentrations of NΔ47-TPH2-R (1.3, 2.5, 5.9, 8.2, and 11.4 mg/mL) were prepared by diluting the stock solution of NΔ47-TPH2-R with L-Phe buffer collected from the filtrate from up-concentration during the protein purification. Measurements (in batch mode) were performed at the German Electron Synchrotron DESY at the P12 EMBL BioSAXS beamline (Extended data table 2). Data processing and analysis were performed using the ATSAS software package version 2.8.2 ^54^, and final plots were made using Matlab (Mathworks Inc.).

### SAXS modelling

10000 different conformations of the N-terminal (residues 46-60) and C-terminal (residues 136-145) of our NMR structure of L-Phe:NΔ47-TPH2-R were generated using RANCH ^55^ with the following parameters: The chain type was set to random coil, the symmetry type to mixture, and the percentage of symmetric conformers to 50%. Subsequently, GAJOE ^55^ was run on the RANCH structure pool using default settings and fitting to the 11.4 mg/mL SAXS curve. To estimate the volume fraction of each conformer, the output structures from GAJOE were submitted to OLIGOMER ^56^ with a constant added as additional component and keeping the remaining parameters as the default settings. In case the error on the volume fraction of a given conformer was larger than the estimated volume fraction, this conformer was taken out of the ensemble from GAJOE, and OLIGOMER was re-run with the reduced set of conformers.

### EM sample preparation and EM data acquisition

The L-Phe sample for cryo-EM (21 μM NΔ47-TPH2, 50 μM L-Phe, 20 mM Hepes, 300 mM (NH_4_)_2_SO_4,_ pH 7.0, and residual components from dilution of MBP-3CP into binding buffer during purification <14 μM DTT, <700 μM Tris, <140 μM EDTA, <0.3% V/V glycerol) was prepared by adding L-Phe from a stock solution to freshly purified NΔ47-TPH2. To prepare the grids, 4 μL of sample were applied onto glow-discharged R2/2 QUANTIFOIL grids and excess sample was removed using an FEI Vitrobot loaded with pre-wet filter paper with a blotting time of 6.5 s, a blotting force of 9, and at 100% humidity and 4 °C. The grid was subsequently flash-frozen by plunging into liquid ethane. Cryo-EM grids were screened, and EM images from the grids with optimal ice thickness and particle distribution were acquired at 300 kV on an FEI Titan Krios electron microscope (Thermo Fisher Scientific, USA), equipped with a Falcon III direct electron detector. Images were recorded via Thermo Fischer EPU 2.1 in integration mode at 132,000x magnification, corresponding to a calibrated pixel size of 1.06 Å at the specimen level, using an exposure time of 1.02 s with 40 movie frames and a total dose of 78 e^−^ per Å^2^. The cryo-EM grids of NΔ47-TPH2 without L-Phe was prepared essentially in the same way, except that the sample contained (18 μM NΔ47-TPH2, 20 mM Hepes, 300 mM (NH_4_)_2_SO_4_, pH 7.0, <14 μM DTT, <700 μM Tris, <140 μM EDTA, <0.3% V/V glycerol). The EM images were acquired at 300kV on an FEI Titan Krios electron microscope (Thermo Fisher Scientific, USA), equipped with a Cs corrector and a Falcon III direct electron detector. Images were recorded via Thermo Fischer EPU 2.1 in integration mode at 153,750x magnification, corresponding to a calibrated pixel size of 1.16 Å at the specimen level, using an exposure time of 1.02 second with 40 movie frames and a total dose of 60 e^−^ per Å^2^. The cryo-EM grids of full-length TPH2 without L-Phe (16 μM TPH2, 20 mM Hepes, 300 mM (NH_4_)_2_SO_4_, pH 7.0, <14 μM DTT, <700 μM Tris, <140 μM EDTA, <0.3% V/V glycerol) was prepared in the same way and recorded with the same settings as NΔ47-TPH2 without L-Phe.

### Image processing

Frames were dose-weighted, aligned, and summed by MotionCor 2.0 ^57^. The defocus values were determined using Gctf ^58^. Summed micrographs were manually evaluated in the COW-MicrographQualityChecker (http://www.cow-em.de). Micrographs with isotropic Thong rings and a defocus range of 1.5 μm to 5 μm that showed clear particles were selected and processed further.

For NΔ47-TPH2 with L-Phe, a total of 1639 summed micrographs were retained for processing, from which about 1.6 million particles were picked using Gautomatch (https://www2.mrc-lmb.cam.ac.uk/research/locally-developed-software/zhang-software/). The picked particles were extracted with a box size of 160×160 pixels and binned to 80 × 80 pixels (pixel size of 2.12 Å) in RELION 3.1 (http://www2.mrc-lmb.cam.ac.uk/relion/index.php/Main_Page). Several rounds of reference-free 2D classification were performed, and classes showing empty boxes, ice/ethane contaminations, or uninterpretable features were removed, yielding 697,589 “good particles”. A subset of 100,000 particles were used to generate an initial 3D map via the *ab initio* reconstruction in cryoSPARC ^59^, without symmetry imposed (i.e. C1 symmetry). The *ab initio* model from cryoSPARC was low-pass filtered to 30 Å resolution and used as the starting reference for subsequent 3D classification in RELION 3.0. The 697,589 “good particles” from 2D classification were 3D classified into eight classes without any mask applied nor symmetry imposed. The 3D classes showing clear density for both CDs and RDs were selected, yielding 310,165 particles. These particles were further classified into five classes without symmetry or mask, and the one class containing 58,616 particles that shows clear RD densities on both sides of the CDs was selected and further refined by Refine3D in RELION 3.1 into a map of the entire NΔ47-TPH2 with an average resolution of 8.9 Å (ranging from ~7 Å of the CDs to ~15 Å of the RDs), without symmetry imposed. To reconstruct a higher resolution map for the CDs, the 697,589 “good particles” from 2D classification were 3D classified into 8 classes with a mask applied around the CDs and TDs (hereafter referred to as the CD tetramer) without symmetry imposed. The best classes showing clear CD density were selected and combined, yielding 241,504 particles, which were further classified into eight classes with a mask applied around the CD tetramer but without symmetry imposed. The best classes containing 197,917 particles were selected, re-extracted with a box size of 360×360 pixels (pixel size of 1.06 Å) and refined with a mask around the CD tetramer and D2 symmetry imposed, resulting in a 4.1 Å map of the CD tetramer. After polishing, the resolution of the CD tetramer map was further improved to 3.9 Å (D2 symmetry imposed). For NΔ47-TPH2 without L-Phe, a total of 1149 summed micrographs were retained, from which 313,195 particles were extracted with a box size of 210×210 pixels and binned to 70 × 70 pixels (pixel size of 3.48 Å) in RELION 3.1 (http://www2.mrc-lmb.cam.ac.uk/relion/index.php/Main_Page). After two rounds of 2D classification, the best 2D class averages showing clear features of TPH2 were selected, containing 131,117 “good particles”. For Full-length TPH2, a total of 628 micrographs were recorded, from which 147,940 particles extracted and subjected to 2D classification. However, the 2D class averages did not show reasonable features of typical TPH2, likely due to the aggregation of particles.

### Model building and refinement

For the model of the CD tetramer, the crystal structure of the *human* TPH2 CD and TD (PDB: 4V06) was used as the template for model building. The crystal structure was rigid body docked into the 3.9 Å EM map of the NΔ47-TPH2 CD tetramer using Chimera v.1.13.1 ^60^. All the side chains were truncated to poly-alanine and the carbon backbone was manually adjusted into the EM density using Coot v. 0.8.9.2 ^61^ with secondary structure restrains. The model was then refined in PHENIX ^62^ via real-space refinement with secondary structure restrains. For the model of the entire NΔ47-TPH2, the refined model of the CD tetramer and the NMR solution structure of the RD dimer obtained in this study were rigid body docked into the EM map of the entire NΔ47-TPH2 and combined without further adjustment or refinement.

## Author contributions

GHJP and IMV conceived the study. GHJP, PH, and BBK supervised the study. BBK and AP designed the NMR experiments and provided input and feedback on NMR data analysis. AP supervised and assisted with the NMR data collection and processing. IMV performed all experiments and data analysis with exception of the cryo-EM data collection and processing. ZZ and IMV prepared cryo-EM samples, and ZZ performed cryo-EM data collection and processing. HS provided feedback on the cryo-EM analysis. NTS performed preliminary cryo-EM experiments. IMV wrote the manuscript apart from the cryo-EM method section, which was written by ZZ. GHJP, BBK, AP, ZZ, and PH provided feed-back and revisions for the manuscript.

## Acknowledgements

The synchrotron SAXS data was collected at beamline P12 operated by EMBL Hamburg at the PETRA III storage ring (DESY, Hamburg, Germany). We would like to thank Melissa Graevert for the assistance in using the beamline. The authors thank Danscatt for funding the SAXS trip. The NMR infrastructure was supported by Novo Nordisk Foundation infrastructure grant [#NNF18OC0032996, cOpenNMR]. The work was in part funded by the Novo Nordisk Foundation Challenge grant REPIN [#NNF18OC0033926; to B.B.K.] and NTS’ Ph.D. study was financially supported by the Independent Research Fund Denmark [DFF-6108-00247; to G.H.J.P.]. The authors would additionally like to thank David F. Nielsen and Signe Sjørup for technical assistance, and Yulian Gavrilov and Alina Kulakova for useful discussions.

## References

1. Udenfriend, S., Clark, C. T. & Titus, E. 5-Hydroxytryptophan Decarboxylase: A New Route of Metabolism of Tryptophan. J. Am. Chem. Soc. 75, 501–502 (1953).

2. Walther, D. J. & Bader, M. A unique central tryptophan hydroxylase isoform. Biochem. Pharmacol. 66, 1673–1680 (2003).

3. Matthes, S., Mosienko, V., Bashammakh, S., Alenina, N. & Bader, M. Tryptophan Hydroxylase as Novel Target for the Treatment of Depressive Disorders. Pharmacology 85, 95–109 (2010).

4. Coates, M. D. et al. Molecular defects in mucosal serotonin content and decreased serotonin reuptake transporter in ulcerative colitis and irritable bowel syndrome. Gastroenterology 126, 1657–1664 (2004).

5. Quednow, B. B., Geyer, M. A. & Halberstadt, A. L. Serotonin and schizophrenia. in Handbook of Behavioral Neuroscience 31, 711–743 (2020).

6. Haghighi, F. et al. Genetic architecture of the human tryptophan hydroxylase 2 Gene: existence of neural isoforms and relevance for major depression. Mol. Psychiatry 13, 813–820 (2008).

7. Fitzpatrick, P. F. Tetrahydropterin-Dependent Amino Acid Hydroxylases. Annu. Rev. Biochem. 68, 355–381 (1999).

8. Carkaci-Salli, N. et al. Functional Domains of Human Tryptophan Hydroxylase 2 (hTPH2). J. Biol. Chem. 281, 28105–28112 (2006).

9. Arturo, E. C. et al. First structure of full-length mammalian phenylalanine hydroxylase reveals the architecture of an autoinhibited tetramer. Proc. Natl. Acad. Sci. 113, 2394–2399 (2016).

10. Yohrling, G. J., Jiang, G. C.-T., Mockus, S. M. & Vrana, K. E. Intersubunit binding domains within tyrosine hydroxylase and tryptophan hydroxylase. J. Neurosci. Res. 61, 313–320 (2000).

11. Vrana, K. E., Walker, S. J., Rucker, P. & Liu, X. A Carboxyl Terminal Leucine Zipper Is Required for Tyrosine Hydroxylase Tetramer Formation. J. Neurochem. 63, 2014–2020 (2002).

12. Fitzpatrick, P. F. The Aromatic Amino Acid Hydroxylases. in Advances in Enzymology and Related Areas of Molecular Biology 74, 235–294 (2000).

13. Murphy, K. L., Zhang, X., Gainetdinov, R. R., Beaulieu, J.-M. & Caron, M. G. A Regulatory Domain in the N Terminus of Tryptophan Hydroxylase 2 Controls Enzyme Expression. J. Biol. Chem. 283, 13216–13224 (2008).

14. Tidemand, K. D. et al. Stabilization of tryptophan hydroxylase 2 by L-phenylalanine-induced dimerization. FEBS Open Bio 1–13 (2016). doi:10.1002/2211-5463.12100

15. Winge, I. et al. Activation and stabilization of human tryptophan hydroxylase 2 by phosphorylation and 14-3-3 binding. Biochem. J. 410, 195–204 (2008).

16. Shiman, R. Relationship between the substrate activation site and catalytic site of phenylalanine hydroxylase. J. Biol. Chem. 255, 10029–10032 (1980).

17. Arturo, E. C. et al. Manipulation of a cation-π sandwich reveals conformational flexibility in phenylalanine hydroxylase. Biochimie 183, 63–77 (2021).

18. Tourian, A. Activation of phenylalanine hydroxylase by phenylalanine. Biochim. Biophys. Acta - Enzymol. 242, 345–354 (1971).

19. Knappskog, P. M., Flatmark, T., Aarden, J. M., Haavik, J. & Martinez, A. Structure/Function Relationships in Human Phenylalanine Hydroxylase. Effect of Terminal Deletions on the Oligomerization, Activation and Cooperativity of Substrate Binding to the Enzyme. Eur. J. Biochem. 242, 813–821 (1996).

20. Meisburger, S. P. et al. Domain movements upon activation of phenylalanine hydroxylase characterized by crystallography and chromatography-coupled small-angle X-ray scattering. J. Am. Chem. Soc. 138, 6506–6516 (2016).

21. Flydal, M. I. et al. Structure of full-length human phenylalanine hydroxylase in complex with tetrahydrobiopterin. Proc. Natl. Acad. Sci. 116, 11229–11234 (2019).

22. Patel, D., Kopec, J., Fitzpatrick, F., McCorvie, T. J. & Yue, W. W. Structural basis for ligand-dependent dimerization of phenylalanine hydroxylase regulatory domain. Sci. Rep. 6, 23748 (2016).

23. Arturo, E. C., Gupta, K., Hansen, M. R., Borne, E. & Jaffe, E. K. Biophysical characterization of full-length human phenylalanine hydroxylase provides a deeper understanding of its quaternary structure equilibrium. J. Biol. Chem. 294, 10131–10145 (2019).

24. Zhang, S., Roberts, K. M. & Fitzpatrick, P. F. Phenylalanine binding is linked to dimerization of the regulatory domain of phenylalanine hydroxylase. Biochemistry 53, 6625–6627 (2014).

25. McKinney, J., Knappskog, P. M. & Haavik, J. Different properties of the central and peripheral forms of human tryptophan hydroxylase. J. Neurochem. 92, 311–320 (2005).

26. Tenner, K., Walther, D. & Bader, M. Influence of human tryptophan hydroxylase 2 N- and C-terminus on enzymatic activity and oligomerization. J. Neurochem. 102, 1887–1894 (2007).

27. Zhang, S., Hinck, C. S. & Fitzpatrick, P. F. The regulatory domain of human tryptophan hydroxylase 1 forms a stable dimer. Biochem. Biophys. Res. Commun. 476, 457–461 (2016).

28. Zhang, S., Huang, T., Ilangovan, U., Hinck, A. P. & Fitzpatrick, P. F. The Solution structure of the Regulatory domain of Tyrosine Hydroxylase. J. Mol. Biol. 1483–1497 (2014). doi:10.1016/j.pestbp.2011.02.012.Investigations

29. Lang, E. J. M., Cross, P. J., Mittelstädt, G., Jameson, G. B. & Parker, E. J. Allosteric ACTion: the varied ACT domains regulating enzymes of amino-acid metabolism. Curr. Opin. Struct. Biol. 29, 102–111 (2014).

30. Cross, P. J., Dobson, R. C. J., Patchett, M. L. & Parker, E. J. Tyrosine Latching of a Regulatory Gate Affords Allosteric Control of Aromatic Amino Acid Biosynthesis. J. Biol. Chem. 286, 10216–10224 (2011).

31. Waudby, C. A., Ramos, A., Cabrita, L. D. & Christodoulou, J. Two-Dimensional NMR Lineshape Analysis. Sci. Rep. 6, 24826 (2016).

32. Skawinska, N. Allosteric regulation of human tryptophan hydroxylase isoform 2 (hTPH2). (PhD thesis, Department of Chemistry, Technical University of Denmark, 2020).

33. Bueno-Carrasco, M. T. et al. Structural mechanism for tyrosine hydroxylase inhibition by dopamine and reactivation by Ser40 phosphorylation. Nat. Commun. 13, 74 (2022).

34. Zhu, K. et al. Cryo-EM Structure and Activator Screening of Human Tryptophan Hydroxylase 2. Front. Pharmacol. 13, (2022).

35. Waisbren, S. E. et al. Improved Measurement of Brain Phenylalanine and Tyrosine Related to Neuropsychological Functioning in Phenylketonuria. in JIMD Reports 4, 77–86 (2016).

36. Möller, H. E., Ullrich, K. & Weglage, J. In vivo proton magnetic resonance spectroscopy in phenylketonuria. Eur. J. Pediatr. 159, S121–S125 (2000).

37. Khan, C. A., Meisburger, S. P., Ando, N. & Fitzpatrick, P. F. The phenylketonuria-associated substitution R68S converts phenylalanine hydroxylase to a constitutively active enzyme but reduces its stability. J. Biol. Chem. 294, 4359–4367 (2019).

38. Jumper, J. et al. Highly accurate protein structure prediction with AlphaFold. Nature 596, 583–589 (2021).

39. de Groot, M. J., Hoeksma, M., Blau, N., Reijngoud, D. J. & van Spronsen, F. J. Pathogenesis of cognitive dysfunction in phenylketonuria: Review of hypotheses. Mol. Genet. Metab. 99, S86–S89 (2010).

40. Winn, S. R. et al. Blood phenylalanine reduction corrects CNS dopamine and serotonin deficiencies and partially improves behavioral performance in adult phenylketonuric mice. Mol. Genet. Metab. 123, 6–20 (2018).

41. Ogawa, S. & Ichinose, H. Effect of metals and phenylalanine on the activity of human tryptophan hydroxylase-2: Comparison with that on tyrosine hydroxylase activity. Neurosci. Lett. 401, 261–265 (2006).

42. Gasteiger, E. et al. Protein Identification and Analysis Tools on the ExPASy Server. in The Proteomics Protocols Handbook 571–607 (Humana Press, 2005). doi:10.1385/1-59259-890-0:571

43. Kazimierczuk, K. & Orekhov, V. Y. Accelerated NMR Spectroscopy by Using Compressed Sensing. Angew. Chemie Int. Ed. 50, 5556–5559 (2011).

44. Delaglio, F. et al. NMRPipe: A multidimensional spectral processing system based on UNIX pipes. J. Biomol. NMR 6, 277–293 (1995).

45. Shen, Y. & Bax, A. Protein backbone and sidechain torsion angles predicted from NMR chemical shifts using artificial neural networks. J. Biomol. NMR 56, 227–241 (2013).

46. Kjaergaard, M., Brander, S. & Poulsen, F. M. Random coil chemical shift for intrinsically disordered proteins: effects of temperature and pH. J. Biomol. NMR 49, 139–149 (2011).

47. Vranken, W. F. et al. The CCPN data model for NMR spectroscopy: Development of a software pipeline. Proteins Struct. Funct. Genet. 59, 687–696 (2005).

48. Coote, P. W. et al. Optimal control theory enables homonuclear decoupling without Bloch–Siegert shifts in NMR spectroscopy. Nat. Commun. 9, 3014 (2018).

49. Güntert, P. & Buchner, L. Combined automated NOE assignment and structure calculation with CYANA. J. Biomol. NMR 62, 453–471 (2015).

50. Schwieters, C. D., Kuszewski, J. J., Tjandra, N. & Clore, G. M. The Xplor-NIH NMR molecular structure determination package. J. Magn. Reson. 160, 65–73 (2003).

51. Schwieters, C. D., Kuszewski, J. J. & Marius Clore, G. Using Xplor-NIH for NMR molecular structure determination. Prog. Nucl. Magn. Reson. Spectrosc. 48, 47–62 (2006).

52. Tian, Y., Schwieters, C. D., Opella, S. J. & Marassi, F. M. A practical implicit solvent potential for NMR structure calculation. J. Magn. Reson. 243, 54–64 (2014).

53. Teilum, K., Kunze, M. B. A., Erlendsson, S. & Kragelund, B. B. (S)Pinning down protein interactions by NMR. Protein Sci. 26, 436–451 (2017).

54. Franke, D. et al. ATSAS 2.8 : a comprehensive data analysis suite for small-angle scattering from macromolecular solutions. J. Appl. Crystallogr. 50, 1212–1225 (2017).

55. Tria, G., Mertens, H. D. T., Kachala, M. & Svergun, D. I. Advanced ensemble modelling of flexible macromolecules using X-ray solution scattering. IUCrJ 2, 207–217 (2015).

56. Konarev, P. V., Volkov, V. V., Sokolova, A. V., Koch, M. H. J. & Svergun, D. I. PRIMUS : a Windows PC-based system for small-angle scattering data analysis. J. Appl. Crystallogr. 36, 1277–1282 (2003).

57. Zheng, S. Q. et al. MotionCor2: anisotropic correction of beam-induced motion for improved cryo-electron microscopy. Nat. Methods 14, 331–332 (2017).

58. Zhang, K. Gctf: Real-time CTF determination and correction. J. Struct. Biol. 193, 1–12 (2016).

59. Punjani, A., Rubinstein, J. L., Fleet, D. J. & Brubaker, M. A. cryoSPARC: algorithms for rapid unsupervised cryo-EM structure determination. Nat. Methods 14, 290–296 (2017).

60. Pettersen, E. F. et al. UCSF Chimera - A visualization system for exploratory research and analysis. J. Comput. Chem. 25, 1605–1612 (2004).

61. Emsley, P., Lohkamp, B., Scott, W. G. & Cowtan, K. Features and development of Coot. Acta Crystallogr. Sect. D Biol. Crystallogr. 66, 486–501 (2010).

62. Afonine, P. V. et al. Towards automated crystallographic structure refinement with phenix.refine. Acta Crystallogr. Sect. D Biol. Crystallogr. 68, 352–367 (2012).

63. Chen, V. B. et al. MolProbity : all-atom structure validation for macromolecular crystallography. Acta Crystallogr. Sect. D Biol. Crystallogr. 66, 12–21 (2010).

64. Trewhella, J. et al. 2017 publication guidelines for structural modelling of small-angle scattering data from biomolecules in solution: an update. Acta Crystallogr. Sect. D Struct. Biol. 73, 710–728 (2017).

65. Sievers, F. et al. Fast, scalable generation of high-quality protein multiple sequence alignments using Clustal Omega. Mol. Syst. Biol. 7, 539 (2011).

