## Extended Data for "Structural characterization of human tryptophan hydroxylase 2 reveals L-Phe as the superior regulatory domain ligand relevant for serotonin biosynthesis"

Extended data figure 1: $^1H$-$^{15}N$ HSQC spectrum of NΔ47-TPH2-R with 10 mM L-Phe, $^1H$-$^{13}C$ HSQC spectrum of 13C-labelled L-Phe with NΔ47-TPH2-R, and secondary structure prediction based on chemical shift values.

Extended data table 1: NMR structural constraints and refinement statistics

Extended data figure 2: $R_1$ and $R_2$ backbone relaxation rates and heteronuclear $^1H$-$^{15}N$ NOE relaxations of NΔ47-TPH2-R with L-Phe.

Extended data figure 3: SAXS modelling fit, modelled SAXS conformers, and SAXS scattering curves.

Extended data table 2: Details on SAXS data collection, data analysis, and modelling.

Extended data figure 4: Panels of a $^{13}C$/$^{15}N$-filtered $^{13}C$-NOESY-HSQC of NΔ47-TPH2-R with L-Phe

Extended data figure 5: Representative spectra from titration of NΔ47-TPH2-R with L-Phe, and TITAN line-shape analysis.

Extended data figure 6: $R_1$ and $R_2$ backbone relaxation rates of NΔ47-TPH2-R with and without L-Phe. Secondary structure propensity and secondary chemical shifts of NΔ47-TPH2-R with and without L-Phe.

Extended data figure 7: Titration data from the titration of $^{13}C$,$^{15}N$-labelled L-Phe with un-labelled NΔ47-TPH2-R.

Extended data table 3: Cryo-EM structural statistics

Extended data figure 8: Structural alignment of tyrosine hydroxylase and tryptophan hydroxylase 2 tetramers. Cryo-EM 2D class averages of full-length TPH2

Extended data figure 9: AlphaFold2 prediction of full-length TPH2

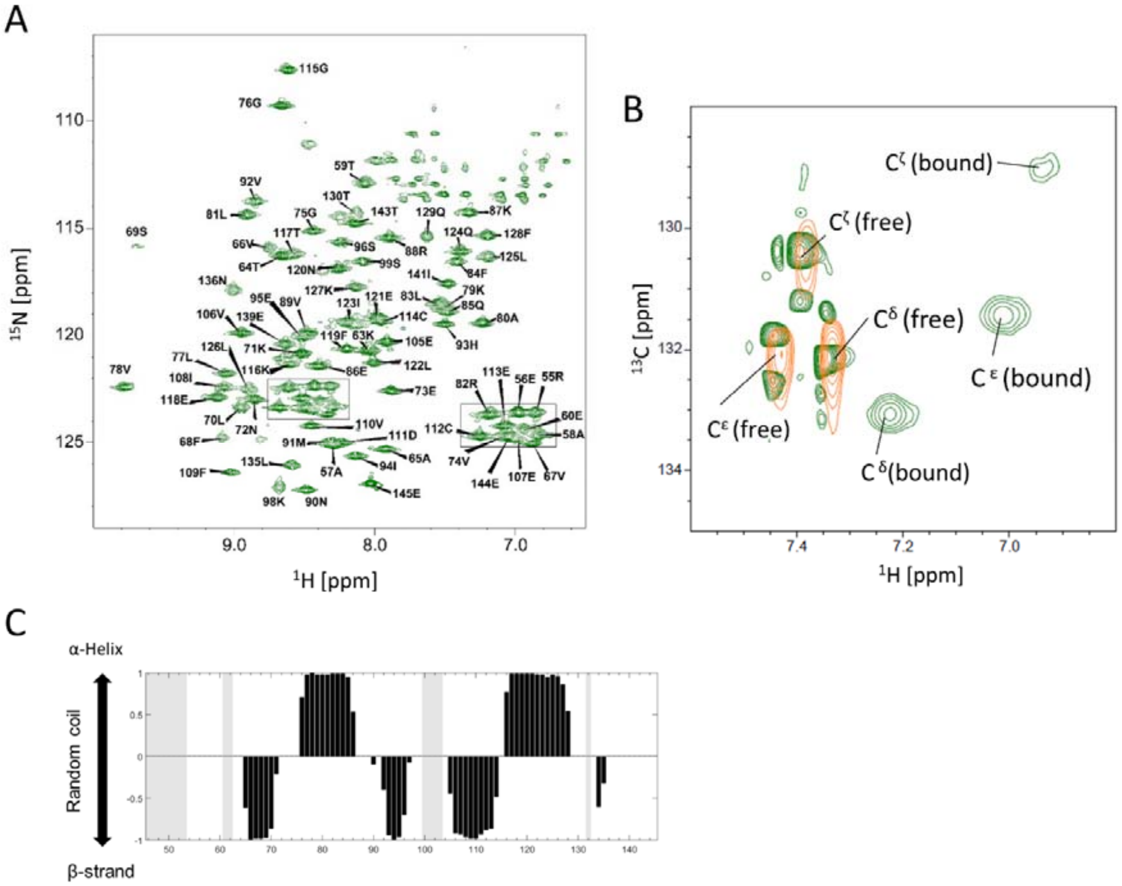

*Extended data figure 1 A) $^{1}$H-$^{15}$N HSQC spectrum of NΔ47-TPH2-R in the presence of 10 mM L-Phe. B) $^{1}$H-$^{13}$C HSQC spectrum of $^{13}$C-labelled L-Phe with NΔ47-TPH2-R (green) and free $^{13}$C-labelled L-Phe (orange). C) Secondary structure prediction based on chemical shift values. Calculated using TALOS-N. Grey areas indicate residues where there is no chemical shift information that allows for calculation of secondary structure propensity.*

*Extended Data Table 1 NMR structural statistics, PDB ID: 7QRI*

| | L-Phe:NΔ47-TPH2-R |
|---|---|
| **NMR distance and dihedral constraints** | |
| Distance constraints (dimer) | |
| Total NOE | 2171 |
| Intra-residue | 554 |
| Inter-residue | 1617 |
| Sequential ($\|i-j\|$ = 1) | 541 |
| Medium-range (1<$\|i-j\|$ < 5) | 348 |
| Long-range ($\|i-j\|$ > 5) | 728 |
| Intermolecular (protein-protein) | 118 |
| Intermolecular (protein-ligand) | 68 |
| Total dihedral angle restraints (monomer) | 108 |
| $\phi$ | 54 |
| $\psi$ | 54 |
| **Structure statistics** | |
| Violations (mean and s.d.) | |
| Distance constraints (Å) | 0.043±0.003 |
| Dihedral angle constraints (º) | 0.53±0.07 |
| Max. dihedral angle violation (º) | 6.17 |
| Max. distance constraint violation (Å) | 0.635 |
| Deviations from idealized geometry | |
| Bond lengths (Å) | 0.006±0.000 |
| Bond angles (º) | 2.29±0.01 |
| Impropers (º) | 2.436±0.006 |
| Average pairwise r.m.s. deviation** (Å) | |
| Protein (All heavy) | 1.3±0.1 |
| Protein (Backbone) | 0.6±0.1 |
| L-Phe ligands (All heavy) | 1.0±0.6 |
| Ramachandran statistics*** | |
| Favoured region | 99% |
| Allowed region | 1% |
| Outliers | 0% |

***Pairwise r.m.s. deviation was calculated among 20 refined structures using residues 64-97 and 105-130 for the protein (protein core defined by PDB server) and including both L-Phe ligands.*

****Ramachandran statistics were obtained from the wwPDB structure validation report [63], which included residues 64-97 and 105-130 in the analysis.*

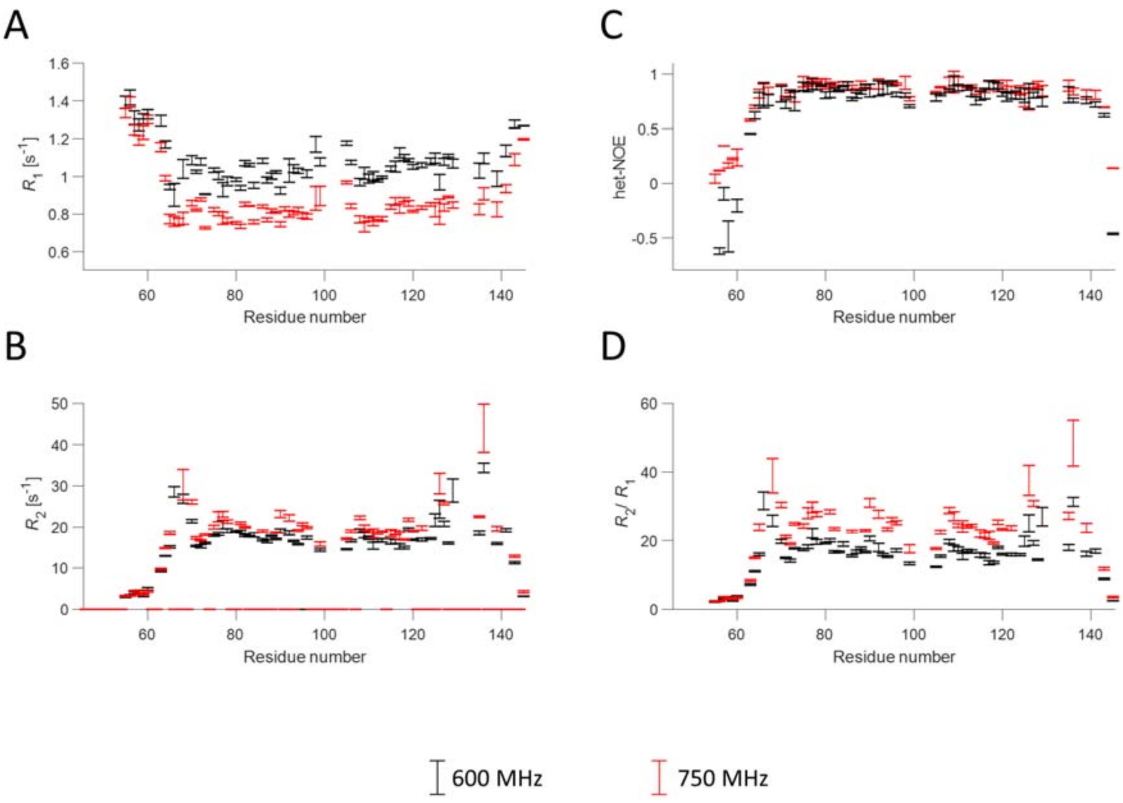

*Extended data figure 2 Backbone $^{15}N$ relaxation parameters of NΔ47-TPH2-R with 10 mM L-Phe at 30 $^{0}C$. A) Longitudinal ($R_1$) relaxation rates, B) Transverse ($R_2$) relaxation rates, C) heteronuclear $^{1}H$-$^{15}N$ NOE relaxations, and D) $R_2/R_1$*

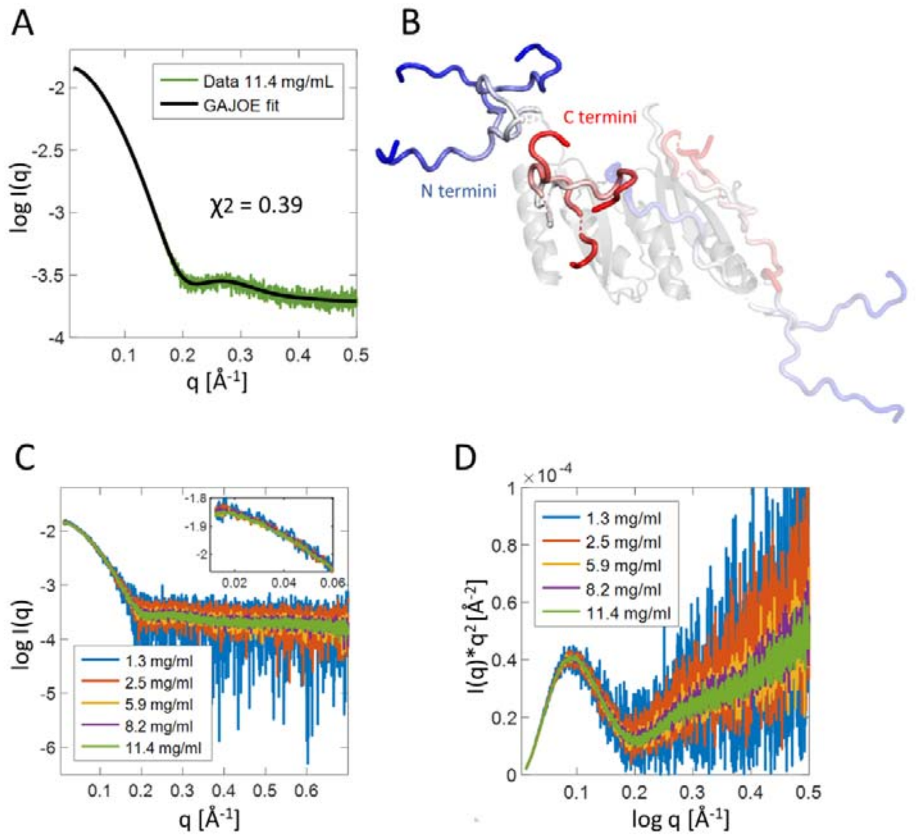

*Extended data figure 3 A) GAJOE [55] fit using a RANCH [55] structure pool of 10000 conformers of residues 46-60 and 136-145, to an 11.4 mg/mL SAXS curve of NΔ47-TPH2-R with L-Phe, resulting in a $\chi^2$ of 0.39. B) Overlay of the three different N- and C-terminal RANCH conformers determined by GAJOE (resulting in the fit in A), illustrating the randomness of the N- and C-termini conformations. C) SAXS curves of NΔ47-TPH2-R in different protein concentrations in the presence of 10 mM L-Phe, showing that NΔ47-TPH2-R is in the same conformation in the range 1.3-11.4 mg/mL. The insert shows a zoom-in of the low q-region revealing no attractive or repulsive interactions in the concentration range. D) Kratky representation of the SAXS curves in C indicating a folded but flexible protein.*

*Extended Data Table 2 SAXS data collection and processing details. The table was prepared according to the guidelines from [64]. C) Details on the samples and the data collection parameters, and the software used for data analysis. D) Structural parameters of NΔ47-TPH2-R calculated based on the SAXS data. F) Details on the SAXS modelling.*

| **A) Sample details** | |
|---|---|
| | NΔ47-TPH2-R |
| Organism | *Homo sapiens* |
| Expression host | E. Coli BL21 |
| Source | Expressed and purified in-house |
| Uniprot sequence ID | Q8IWU9, reidues 48-145 + additional N-terminal Gly-Pro from MBP purification tag |
| Extinction coefficient ($A_{280}$, $M^{-1}cm^{-1}$) | 5500 |
| Molecular mass *M* from chemical composition (kDa) | 11.266 |
| Sample concentration range (mg/mL) | 1.3-11.4 |
| Solvent composition | 12 mM $Na_2HPO_4$/8 mM $NaH_2PO_4$, 100 mM $(NH_4)_2SO_4$, 10 mM L-Phe pH 7.0 (adjusted with $NH_4OH$) |

| **B) SAXS data collection parameters** | |
|---|---|
| Instrument | P12 BioSAXS beamline (PETRAIII) |
| Date | Dec-2019 |
| Detector | Pilatus6m |
| Wavelength (Å) | 1.23984 |
| Beam size ($mm^2$) | At the detector: 0.2x0.12 |
| Detector distance (m) | 3.0 |
| *q*-measurement range ($nm^{-1}$) | 0.03-7.29 |
| Absolute scaling method | Comparison with scattering from pure $H_2O$ |
| Normalization | To transmitted intensity by beam-stop counter |
| Monitoring for radiation damage | Frame-by-frame comparison |
| Exposure time (s) | 0.045 |
| Number of frames | 20 |
| Sample configuration | Quartz glass capillary |
| Sample temperature (ºC) | 20 |

| **C) Software employed for SAXS data reduction, analysis, and interpretation** | |
|---|---|
| SAS data reduction | Primusqt from Atsas [54] |
| Calculation of ε from sequence | ProtParam from ExPaSy [42] |
| Atomic structure modelling | EOM from Atsas [55] |
| Molecular graphics | PyMOL 2.3.3 (Schrödinger) |

| **D) Structural Parameters** | | | | | |
|---|---|---|---|---|---|
| Sample concentration (mg/mL) | 1.3 | 2.5 | 5.6 | 8.2 | 11.4 |
| | | | Guinier Analysis | | |
| *I(0)/c* | 0.015 | 0.015 | 0.015 | 0.015 | 0.015 |
| $R_g$ (Å) | 20.9±0.5 | 20.9±0.3 | 20.7±0.3 | 20.8±0.1 | 20.6±0.1 |
| *q*-range ($Å^{-1}$) | 0.01470-0.06234 | 0.01221-0.06234 | 0.01526-0.06234 | 0.01277-0.06261 | 0.01498-0.06261 |
| Fidelity (%) | 80 | 96 | 98 | 99 | 99 |
| *M* from *I(0) (kDa)* | 21 | 21 | 21 | 21 | 21 |
| | | | *P(r)* analysis | | |
| *I(0)/c* | 0.0151 ±0.0001 | 0.01484 ±0.00005 | 0.01470 ±0.00004 | 0.01484 ±0.00003 | 0.01462 ±0.0003 |
| $R_g$ (Å) | 21.2±0.2 | 21.1±0.1 | 20.85±0.07 | 20.99±0.05 | 20.77±0.05 |
| $d_{max}$ (Å) | 67.5 | 70.0 | 68.9 | 69.5 | 69.7 |
| *q*-range ($Å^{-1}$) | 0.01443-0.39160 | 0.01194-0.39326 | 0.01498-0.39326 | 0.01249-0.39326 | 0.01470-0.43756 |
| Quality estimate (%) | 81 | 79 | 75 | 79 | 76 |
| *M* from *I(0) (kDa)* | 21 | 21 | 20 | 21 | 20 |
| *Porod Volume ($Å^3$)* | 48380 | 50300 | 49440 | 51830 | 50780 |

| **F) Atomistic modelling** | |
|---|---|
| | 11.4 mg/mL NΔ47-TPH2-R |
| Method | RANCH + GAJOE (Ensemble) |
| *q*-range for fitting ($Å^{-1}$) | 0.01-0.5 |
| Symmetry | P2 |
| $\chi^2$ | 0.39 |
| Constant adjustment to intensities | 0.001 |

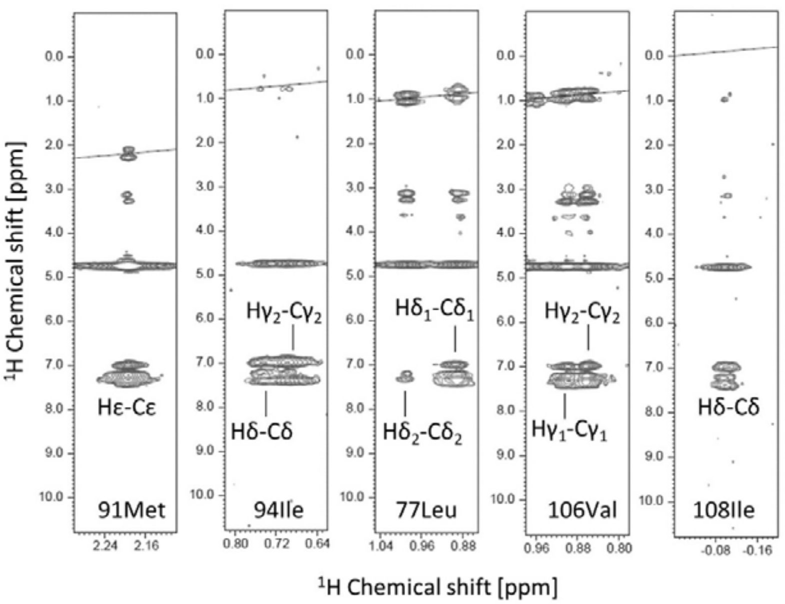

*Extended data figure 4 Panels of a $^{13}C/^{15}N$-filtered $^{13}C$-NOESY-HSQC of NΔ47-TPH2-R with cross-peaks to the L-Phe ligands.*

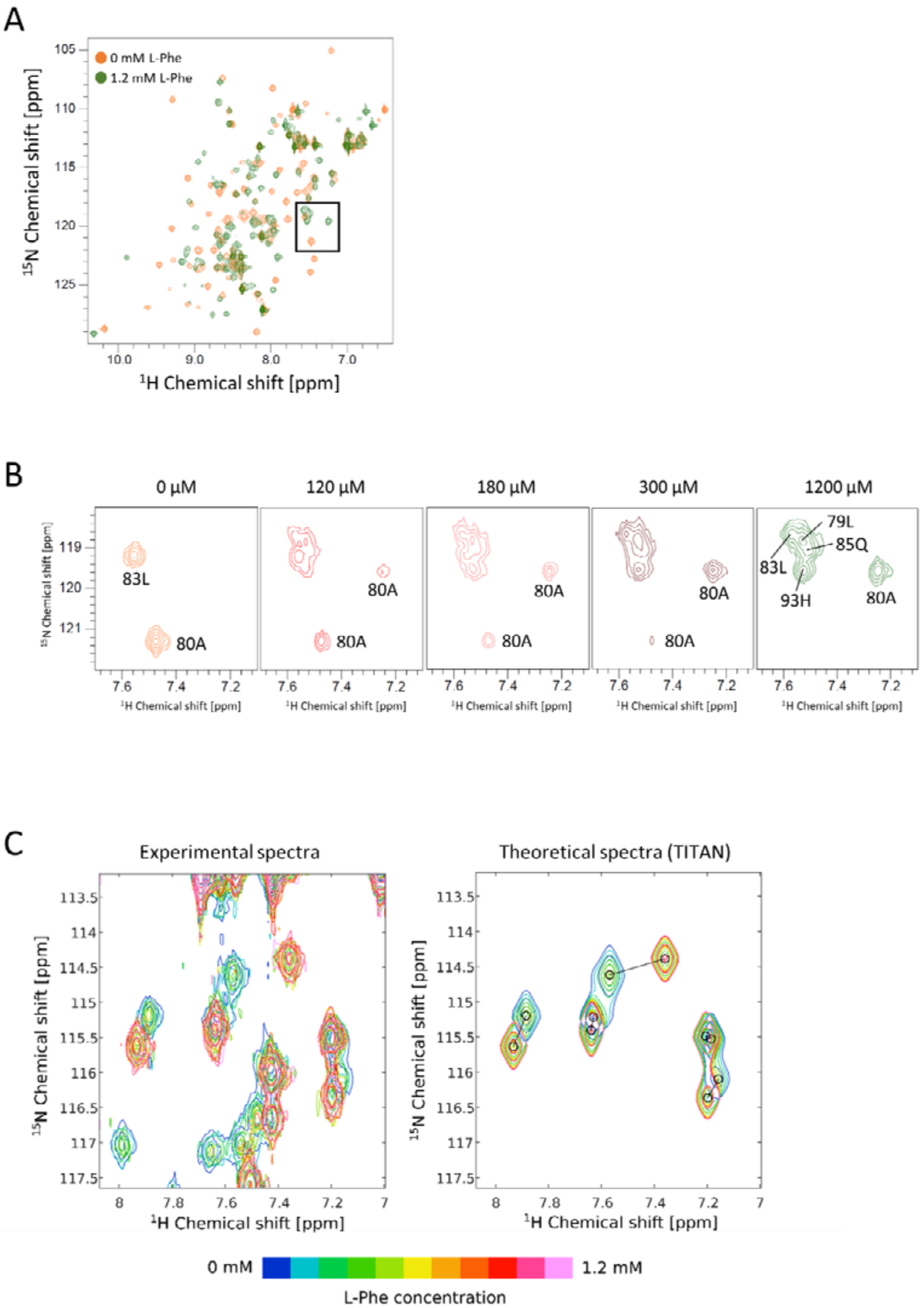

*Extended data figure 5 A) $^{1}H$-$^{15}N$ HSQC spectra of NΔ47-TPH2-R with 0 mM L-Phe (orange) and 1 mM L-Phe (green). B) zoom in of the area marked by a black square in A, showing selected spectra with different L-Phe concentrations illustrating the slow exchange kinetics of the system. C) Side-by-side comparison from the titration analysis with TITAN showing a selected region of experimental (left) and theoretical (right) $^{1}H$-$^{15}N$ HSQC spectra of 200 µM NΔ47-TPH2-R with 0-1 mM L-Phe.*

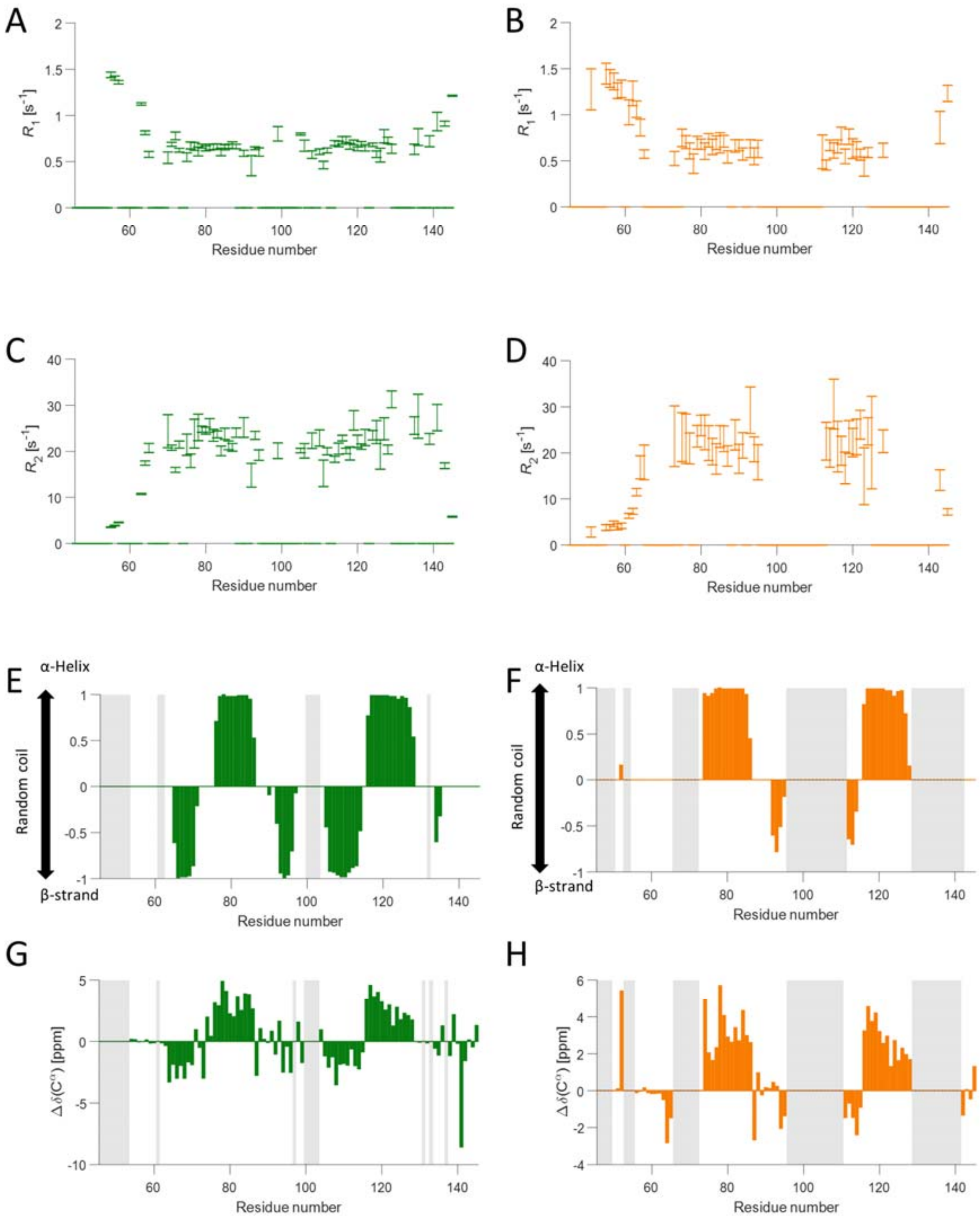

*Extended data figure 6 A-D Backbone $^{15}N$ relaxation parameters of free and L-Phe bound NΔ47-TPH2-R at 20 $^{0}C$ recorded on a 750 MHz spectrometer. Longitudinal ($R_1$) relaxation rates of A) L-Phe bound NΔ47-TPH2-R and B) Free NΔ47-TPH2-R. Transverse ($R_2$) relaxation rates of C) L-Phe bound NΔ47-TPH2-R and D) Free NΔ47-TPH2-R. Secondary structure propensity calculated by TALOS-N of E) L-Phe bound NΔ47-TPH2-R and F) free NΔ47-TPH2-R. Secondary chemical shifts of G) L-Phe bound NΔ47-TPH2-R and H) free NΔ47-TPH2-R. Grey areas in E, F, G, and H indicate residues where chemical shift information that allows for calculation of the parameters are missing.*

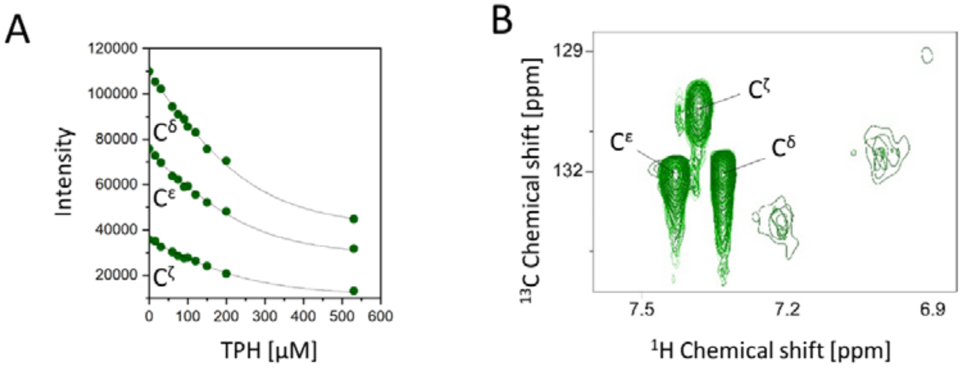

*Extended data figure 7 A) Titration data from the titration of $^{13}C$,$^{15}N$-labelled L-Phe with un-labeled NΔ47-TPH2-R. The intensity decays of the 3 peaks (shown in B) are plotted as green circles. The fits are shown as solid lines. B) $^{1}H$-$^{13}C$ aromatic HSQC spectra of $^{13}C$-labeled L-Phe and different concentrations of un-labeled NΔ47-TPH2-R.*

*Extended Data Table 3* Cryo-EM structural statistics

| | Catalytic core of NΔ47-TPH2 with 50 μM L-Phe (EMD-15853) (PDB 8B4P) | Complete map of NΔ47-TPH2 with 50 μM L-Phe (EMD-15852) (PDB 8B4P) |
|---|---|---|
| Data collection and processing | | |
| Magnification | 120,700 | 120,700 |
| Voltage (kV) | 300 | 300 |
| Electron exposure ($e^{-}/Å^2$) | 72 | 72 |
| Defocus range (μm) | 1-4 | 1-4 |
| Pixel size (Å) | 1.16 | 1.16 |
| Symmetry imposed | C1 | C1 |
| Initial particle images (no.) | 1.6 million | 1.6 million |
| Final particle images (no.) | 197,917 | 58,616 |
| Map resolution (Å) | 3.9 | 8.9 |
| FSC threshold | 0.143 | 0.143 |
| Map resolution range (Å) | | |
| Catalytic domain | 3.9 | 6-9 |
| Regulatory domain | --- | 15-20 |
| Refinement | | |
| Initial model used (PDB code) | 4V06 | 4V06 |
| Model resolution (Å) | 4.8 | --- |
| FSC threshold | 0.5 | |
| Model resolution range (Å) | --- | --- |
| Map sharpening *B* factor ($Å^2$) | -250 | -300 |
| Model composition | | |
| Non-hydrogen atoms | 6873 | |
| Protein residues | 1362 | |
| Ligands | 0 | |
| *B* factors ($Å^2$) | | |
| Protein | --- | --- |
| Ligand | --- | --- |
| R.m.s. deviations | | |
| Bond lengths (Å) | 0.005 | --- |
| Bond angles (°) | 1.17 | --- |
| Validation | | |
| MolProbity score | 0.78 | --- |
| Clashscore | 0.49 | --- |
| Poor rotamers (%) | 0 | --- |
| Ramachandran plot | | |
| Favored (%) | 97.42 | --- |
| Allowed (%) | 2.58 | --- |
| Disallowed (%) | 0 | --- |

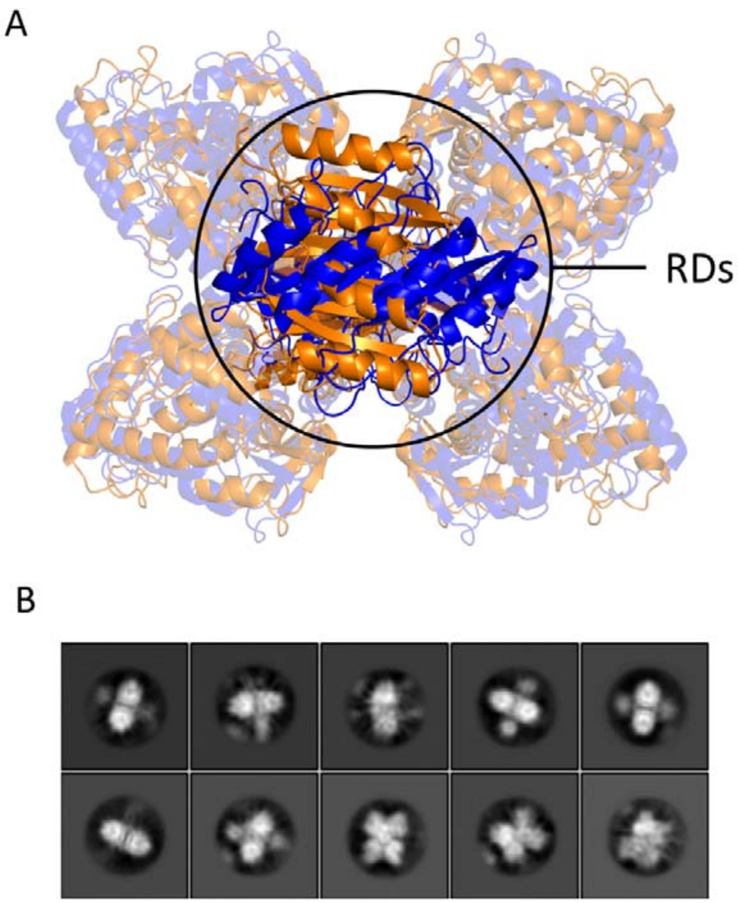

*Extended data figure 8 A) Cryo-EM structure of tyrosine hydroxylase (blue, 6ZVP [33]) aligned with the cryo-EM structure of TPH2 from this study (orange, rigid body model), showing the different orientation of the RD dimers in the tetramers. B) 2D class averages from grids with full-length TPH2.*

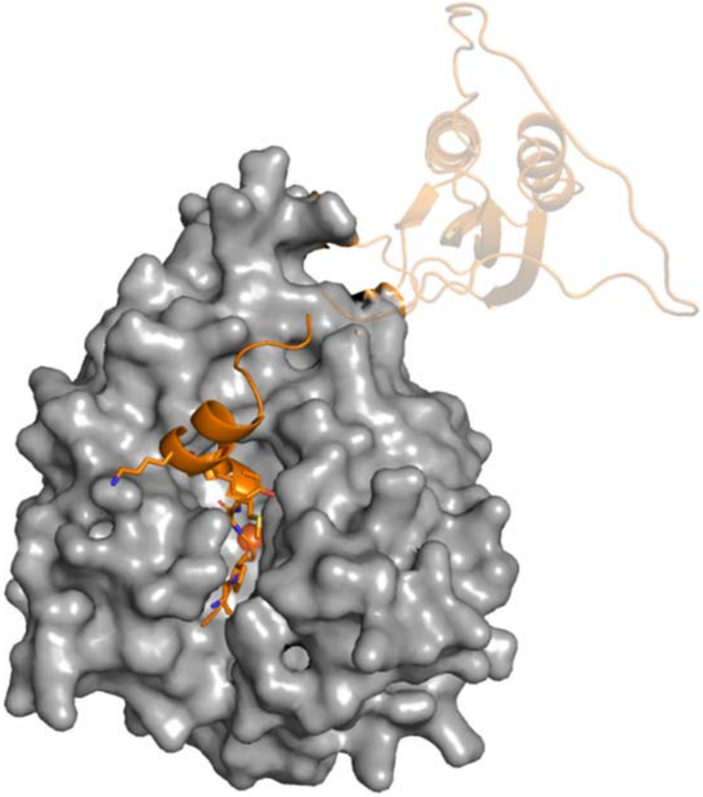

*Extended data figure 9 Structure prediction of full-length TPH2 from alphaFold2 [38]. The catalytic domain is shown as a grey surface, residues 1-21 as orange cartoon, and the rest of the regulatory domain as faded orange cartoon.*
